# Emergent multilevel selection in a simple spatial model of the evolution of altruism

**DOI:** 10.1101/2021.11.30.469839

**Authors:** Rutger Hermsen

## Abstract

Theories on the evolutionary origins of altruistic behavior have a long history and have become a canonical part of the theory of evolution. Nevertheless, the mechanisms that allow altruism to appear and persist are still incompletely understood. It is well known, however, that the spatial structure of populations is an important determinant. In both theoretical and experimental studies, much attention has been devoted to populations that are subdivided into discrete groups. Such studies typically imposed the structure and dynamics of the groups by hand. Here, we instead present a simple individual-based model in which altruistic organisms spontaneously self-organize into spatially separated colonies that themselves reproduce by binary fission and hence behave as Darwinian entities in their own right. Using software to automatically track the rise and fall of colonies, we are able to apply formal theory on multilevel selection and thus quantify the within- and among-group dynamics. This reveals that individual colonies inevitably succumb to defectors in a within-colony “tragedy of the commons”. Even so, altruism persists in the population because more altruistic colonies reproduce more frequently and drive less altruistic ones to extinction. Evidently, the colonies promote the selection of altruism but in turn depend on altruism for their existence; the selection of altruism hence involves a kind of evolutionary bootstrapping. The emergence of the colonies also depends crucially on the length scales of motility, altruism, and competition. This reconfirms the general relevance of these scales for social evolution, but also stresses that their impact can only be understood fully in the light of the emergent eco-evolutionary spatial patterns. The results also suggest that emergent spatial population patterns can function as a starting point for transitions of individuality.

## 1 Introduction

Over the past decades, a rich body of theoretical research has been devoted to the evolution of social behaviors [1, 2]. In particular, much theory has focused on the evolution of cooperation [3, 4], and more narrowly, altruism [5, 6]: behavior that is costly to the actor but beneficial to its interaction partners. Historically, how natural selection could favor altruism has been a puzzle, but in broad terms the solution has long been understood: altruism can be selected if its benefits accrue disproportionately to altruists, thus offsetting their costs [3, 7–9]. Nevertheless, the mechanisms that allow such an interaction structure to arise and persist are still a matter of intense study and debate [4].

Many classical studies considered populations that are subdivided into distinct groups (*e.g*., [10–14]). In such a group structure, altruistic behavior can be selected provided altruists tend to be grouped together and groups with a higher proportion of altruists tend to have higher mean fitness [11]. In nearly all theoretical models of group selection, the group structure and group-level dynamics are imposed or presupposed by the definition of the model. In contrast, we here present a very simple individual-based model in which altruistic organisms *self-organize* into discrete colonies. Moreover, these colonies themselves spontaneously reproduce by growth and binary fission and hence act as Darwinian entities in their own right. In time, each individual colony is fated to collapse; but when it does, another colony grows and divides, giving rise to the kind of multi-level dynamics that in most previous models had to be imposed by hand [14–16]. Such rudimentary, emergent higher-level entities could be a first step towards a full “transition of individuality” [17].

The model describes a spatial environment inhabited by motile organisms that reproduce and interact locally. This it has in common with a significant class of earlier models that use various modeling formalism and have been analyzed using a variety of techniques (e.g., [15, 18–22]). In such models, evolutionary and ecological processes become intimately intertwined because the evolutionary processes are strongly affected by spatial self-structuring, while the self-structuring in turn depend on evolved traits.

Both group-structured and spatially explicit models have demonstrated that local interactions combined with local mating and reproduction can foster altruism if motility is limited, because this allows altruists to aggregate in assorted groups or neighborhoods where they mainly benefit each other [19, 23–25]. However, such models have also revealed important limitations [26–29]. If not only social interactions but also competitive interactions take place locally (“soft” selection [30, 31]), altruists in assorted groups or domains tend to compete with other altruists, in which case the benefits of altruism may be largely or fully canceled by the concomitant increased competition. This local Malthusian trap is alleviated somewhat if the local carrying capacity increases with the proportion of altruists (“elastic” selection), which allows clusters of altruists to become net population sources [32]. Importantly, it can also be avoided if competitive interactions reach beyond the social group or neighborhood, so that altruists in assorted clusters can support each other at the expense of others [24, 26, 29, 33]. This highlights the importance of the relative *scales* of motility, altruism, and competition [1]. As a rule, altruism is favored by limited motility and local social interactions, but global competition.

Long-range competition can come in many implicit forms. For instance, the life cycle of organisms may include a dispersal stage such that individuals can first cooperate with relatives and then compete with non-relatives [26]. Alternatively, the group-level dynamics may include a global mixing stage in which groups or neighborhoods are periodically fragmented and new ones are seeded [34–36]; or groups may occasionally go extinct, after which propagules derived from preexisting groups compete to recolonize their spot [16, 37]. To study the effects of the scales of motility, altruism and competition systematically, the model presented here is deliberately designed such that these scales can be set explicitly and independently. As it turns out, their role is much more intricate than anticipated because they play an essential role in the emergence of the colonies and hence in the resulting multilevel ec*σ*evolutionary dynamics.

To quantitatively analyze model simulations, we use software that automatically tracks the rise and fall of colonies. Subsequently, we apply existing formal theory to quantify the contributions to selection at the individual and colony levels. This demonstrates that, within colonies, natural selection favors defectors who profit from the altruists in their neighborhood but do not share in the costs. But colonies characterized by a higher average level of altruism survive longer and reproduce more frequently, resulting in positive selection at the colony level. The steady level of altruism that eventually establishes can be understood as a balance between these forces: a perpetual “tragedy of the commons” [38] within colonies, compensated by positive selection among them.

## 2 Results

### 2.1 Brief description of the model

We start with a brief specification of the model; details are supplied in the Methods.

The model considers a population of discrete individual in a two-dimensional (2D) or one-dimensional (1D) habitat (see Fig. 1a). Individuals possess just one continuous trait *ϕ*, representing their investment in altruistic behavior, and they do only three things: move, in an unbiased fashion modeled by diffusion; die, at a constant (Poisson) rate; and reproduce asexually.

**Figure 1.**
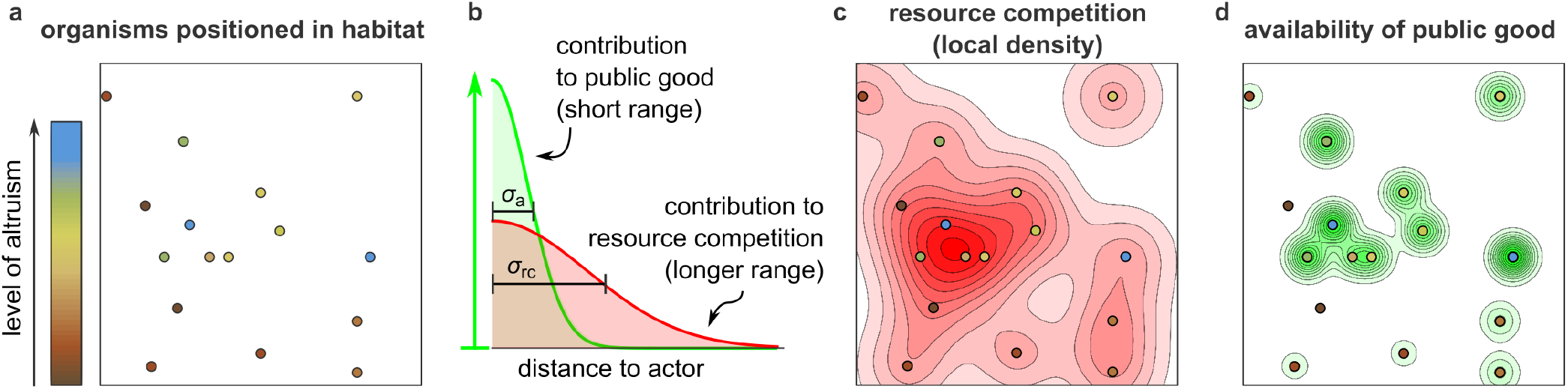
Illustration of the model. (a) The model considers a population of individuals, here represented as circles, in an explicitly spatial habitat. Individuals reproduce, die, and move stochastically, and are characterized by their level of altruism, indicated in color. Altruism is costly to the actor, but beneficial to recipients: it increases their reproduction rate. (b) For concreteness, imagine that each altruist produces a public good and secretes it locally. The contribution of a particular altruist to the concentration of public good falls with the distance (green curve) and increases with the level of altruism of the actor (vertical arrow). At the same time, individuals compete for a limiting resource: the reproduction rate of each individual is inhibited by each individual in its neighborhood (red curve). The scale of altruism *σ*_a_ and the scale of resource competition *σ*_rc_ are indicated. (c & d) The competition experienced at any coordinate (panel c, red contour plot) and the availability of public good at any position (panel d, green contour plot) are obtained by summing up the contributions of all individuals.

The rate of reproduction of each individual depends on three quantities. First, it decreases with the individual’s own investment in altruism: altruism is costly. A level of altruism of *ϕ* = 0.05 means that the individual sacrifices 5% of its reproduction rate relative to a defector (*ϕ* = 0) under the same conditions. Second, the reproduction rate also decreases with the population density in the individual’s local neighborhood. This models competition for resources and establishes a finite carrying capacity. The local population density is measured as a Kernel Density Estimate (KDE), using a normal distribution with standard deviation *σ*_rc_ as the kernel function. This means that individuals compete strongly with each other only if their spatial separation is of order *σ*_rc_ or less (see Fig. 1b, red line, and 1c), so that *σ*_rc_ can be interpreted as the *scale of competition*. Third, an individual’s reproduction rate increases if altruists are present in its local neighborhood (Fig. 1b, green line, and Fig. 1d). The altruism experienced at a given position *y*, denoted *A*(*y*), is again quantified as a KDE, but now individuals are weighted in proportion with their level of altruism *ϕ*. Although the model is not intended to mimic a specific altruistic behavior or mechanism, it is convenient to think of *A*(*y*) as the concentration of some public good secreted by altruistic organisms. If more and more public good is added to the local neighborhood, the benefit eventually saturates. The standard deviation of the kernel function used to calculate *A*(*y*) is called *σ*_a_ and generally differs from *σ*_rc_. Because individuals profit significantly from the public good produced by others only if their separation is of order *σ*_a_ or less, *σ*_a_ can be interpreted as the *scale of altruism*.

It is worth emphasizing that, contrary to some other models [20, 22], complete defectors (with *ϕ* = 0) are perfectly viable; altruism is not required for the survival of the population.

When an individual reproduces, the offspring appears at the coordinates of the parent; afterwards, parent and offspring move independently and thus part ways. Offspring usually inherit the trait value of the parent, but with a small probability a mutation occurs that increases or decreases it at random.

In simulations, space and time are discretized, and periodic boundary conditions are imposed. Default parameter values are listed in Table 1. Throughout the text, the time unit is the inverse of the death rate, called the “generation time”. Importantly, the scale of altruism *σ*_a_ is used as the unit of length and hence *σ*_a_ = 1 by definition. Thus, just two length scales remain: the scale of competition *σ*_rc_ and the scale of motility, *σ*_m_. The latter is defined as the typical (that is, root-mean-square) distance traveled by an individual in a generation time (see Methods).

**Table 1.** Model parameters and their default values for simulations with the 1D and 2D habitat. Units of time, length, and trait are defined such that the death rate *d*, the scale of altruism *σ*_a_, and the reproductive deficit per unit of the trait are 1. All other parameters are expressed in these units.

| parameter | symbol | default (2D) | default (1D) |
| --- | --- | --- | --- |
| death rate | $d$ | 1 | 1 |
| scale of altruism | $\sigma_a$ | 1 | 1 |
| reproductive deficit per unit of the trait value | $c$ | 1 | 1 |
| basal reproduction rate | $g_0$ | 5 | 5 |
| scale of competition | $\sigma_{rc}$ | 4 | 4 |
| motility (diffusion) constant | $k_D$ | $4 \times 10^{-2}$ | $3 \times 10^{-2}$ |
| factor scaling carrying capacity | $K$ | 40 | 100 |
| basal benefit of altruism | $b_0$ | 1 | 0.5 |
| maximal benefit altruism | $b_{\max}$ | 5 | 2 |
| mutation probability upon reproduction | $\mu$ | $1 \times 10^{-3}$ | $5 \times 10^{-4}$ |
| mean effect size of mutations | $m$ | $5 \times 10^{-3}$ | $5 \times 10^{-3}$ |
| scale of motility | $\sigma_m = \sqrt{2k_D/d}$ | 0.283 | 0.245 |

### 2.2 Emergent colonies and multilevel dynamics

The complex behavior of the simple model is illustrated in Fig. 2, which presents results of a single simulation run using a 2D habitat. These results are representative for the default parameters (see replicates in Fig. S1) but the parameters themselves have been chosen deliberately to enable the evolution of altruism. In particular, motility is slow and the scale of competition *σ*_rc_ is four times larger than the scale of altruism *σ*_a_.

**Figure 2.**
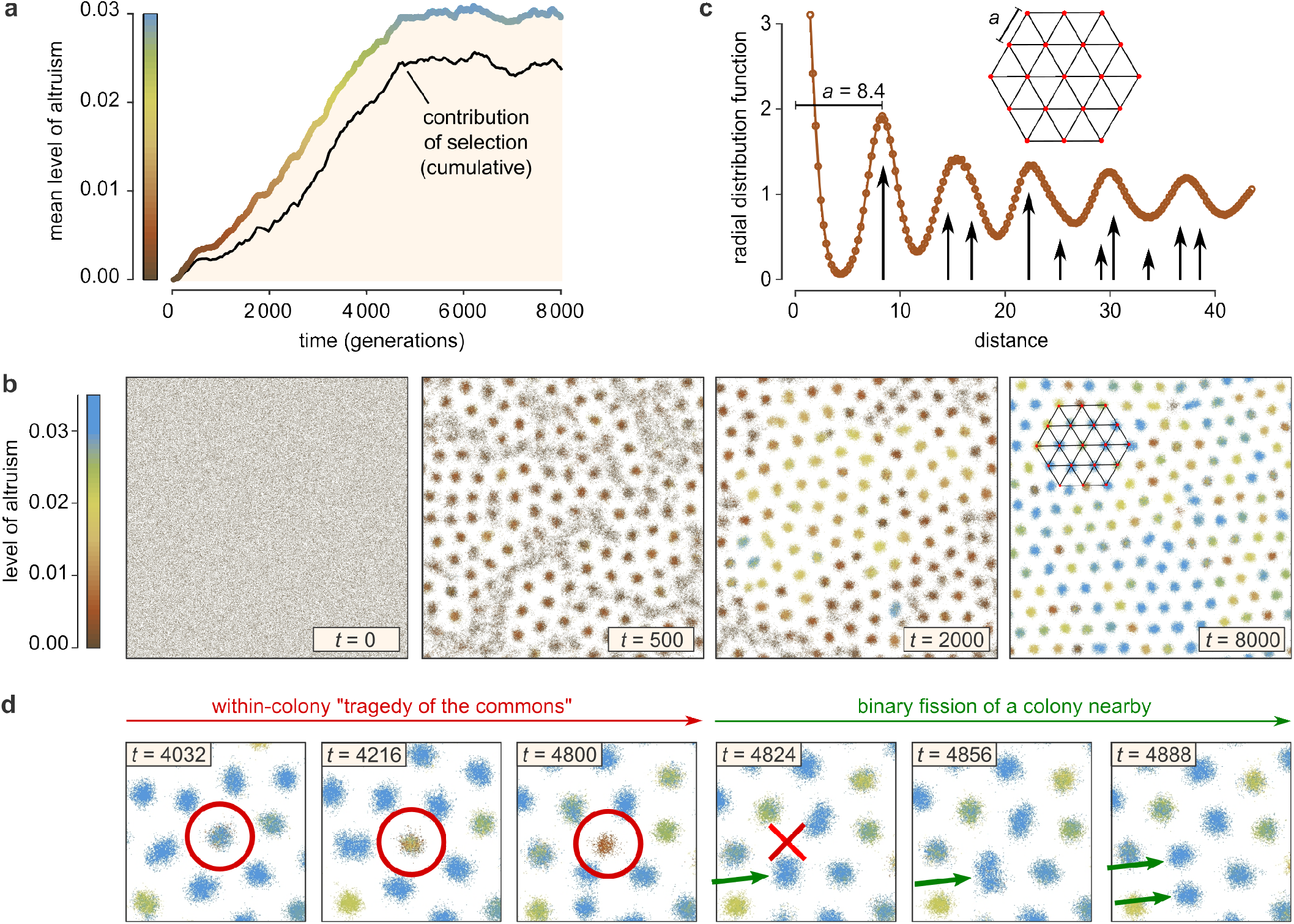
Altruism and colonies emerge in the two-dimensional habitat. Results are shown from a representative simulation run (see Fig. S1 for replicates) with default parameters (see Table 1). (a) Mean level of altruism versus time (thick colored line) as well as the cumulative contribution of natural selection (black), which is consistently positive (see Appendix A.2). (b) Snapshots of the simulation habitat; also see Movies S1–3. In time, the population self-organizes into a hexagonal pattern of discrete colonies. A section of a hexagonal grid is superimposed in the right-most panel. (c) The hexagonal pattern is also apparent from the radial distribution function at *t* = 8000, the distribution of distances between pairs of individuals normalized by the random expectation. Black arrows indicate the distances occurring in an exact hexagonal grid with grid constant *a* = 8.4 and their relative frequency. (d) Enlargements of a small domain of the habitat, showing that the colonies behave like Darwinian entities: they disappear as a result of a within-colony tragedy of the commons [38] (red circle and cross), and reproduce by binary fission (green arrows).

As shown in Fig. 2a, all individuals are initialized as defectors, but in time the mean level of altruism steadily increases (thick colored line) before reaching a plateau. To confirm that this rise is largely due to natural selection rather than random drift or mutational bias, we measured the cumulative contribution of natural selection (black line), which is consistently positive (also see Fig. S1 and Appendix A.2).

The surprising spatial dynamics of the simulation are visualized in the snapshots of Fig. 2b and, more vividly, in Movies S1-3. While individuals are initially distributed uniformly at random, they spontaneously organize into dense colonies surrounded by “exclusion zones”. These colonies subsequently organize into a hexagonal pattern; to illustrate this, a hexagonal grid is overlaid in the right-most panel of Fig. 2b. To further characterize the pattern we determined the radial distribution function, which is defined as the distribution of distances between all pairs of individuals, normalized by the random expectation (Fig. 2c). The long-ranged oscillations in this distribution reveal a lattice constant of *a* ≈ 8.4, consistent with estimates based on the number of colonies found in the habitat (see Methods). The mechanism producing the pattern is analogous to that of the famous Turing patterns in reaction–diffusion systems [39], as we will discuss in some detail below and in Appendix B.

But the pattern is not static. In Fig. 2d, enlargements are shown of a small region of the habitat. Consider the colony marked by the red circle. Initially, the colony is mostly blue, indicating that most individuals in this colony are highly altruistic. In time, however, the color degrades from blue to brown, reflecting a decline in altruism, and eventually the colony goes extinct. As best seen in Movies S1–3, this fate is bestowed on many colonies in the simulation. This suggests that altruistic colonies are sensitive to corruption by defectors that occasionally appear by mutation or invasion from neighboring colonies, resulting in a within-colony “tragedy of the commons” [38]. In the same figure, however, green arrows point to what happens after a colony disappears: a different colony nearby initially grows in size and then spontaneously divides in two, locally restoring the hexagonal pattern. Daughter colonies inherit their over-all color from their parent colony. Importantly, it appears in Movie S1–3 that colonies with a high mean level of altruism divide particularly rapidly and thus manage to multiply and spread.

All in all, these observations suggest that the colonies themselves behave like Darwinian entities: they reproduce by binary fission and show heritable variation in their level of altruism. Moreover, in view of the tragedy of the commons seen within colonies, the colony-level dynamics appear crucial for the evolution of altruism.

### 2.3 Colony formation is crucial for the evolution of altruism

So far, the evidence presented on the dynamics of colonies has been anecdotal and qualitative. To further study the behavior of the model and obtain quantitative results, we now shift to a one-dimensional habitat. Simulations with a one-dimensional habitat are considerably faster, allowing many parameter settings to be explored, and are analyzed more readily, both mathematically and computationally.

#### Automated multilevel lineage tracking

Qualitatively, the behavior of the 1D model is analogous to that of the 2D model. Fig. 3a shows a section of the space-time arena for a simulation with default parameters (see Table 1). The left-hand side of the figure (gray scale) presents the population density. The striped pattern clearly reveals the formation of regularly spaced colonies that can persist for thousands of generations. An algorithm was used to detect these colonies automatically and track them in time (see Methods). The right-hand side of the figure plots the center of mass of the tracked colonies; colors represent the mean level of altruism of the individuals populating the colonies. In the middle part of the figure, density and traces overlap to showcase their consistency. Some traces suddenly end, indicating that the colony went extinct. Such events are detected automatically and indicated with a black square. From the figure, it is apparent that prior to the death of a colony the mean level of altruism always declines, suggesting a tragedy of the commons. In other places, traces suddenly fork, which is also automatically marked, this time with orange circles. Clearly, the colonies in the 1D habitat reproduce by binary fission (like their 2D counterparts); the daughter colonies inherit their color from their parent. Again it appears that more altruistic colonies divide more frequently.

**Figure 3.**
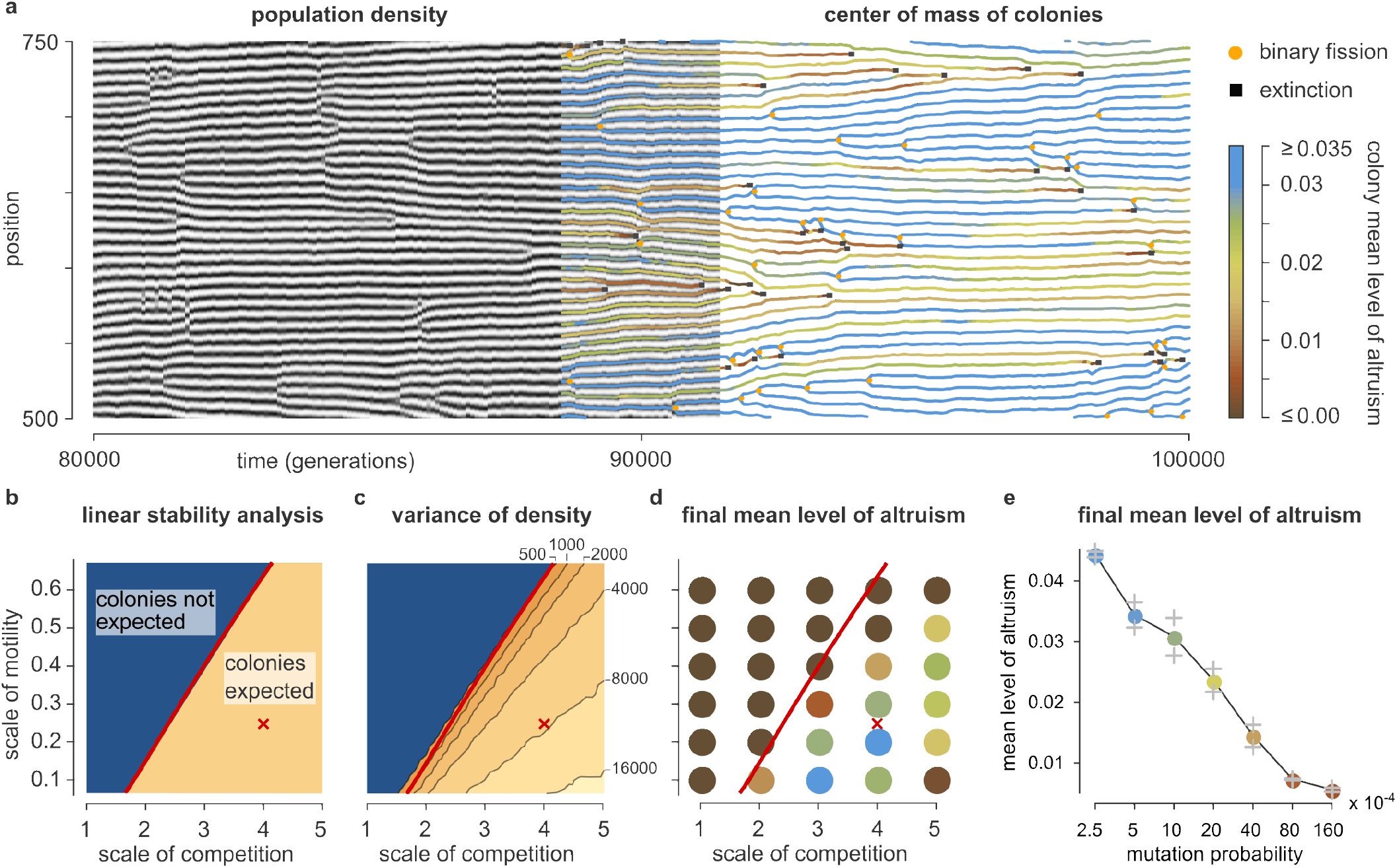
The origins of altruism and colony formation in the one-dimensional habitat. (a) Dynamics of a representative simulation run (see Fig. S4) with default parameters (Table 1). A small domain of space-time is visualized. The left-hand part of the figure shows the local population density; the striped pattern indicates that regularly spaced colonies develop. The right-hand side plots the center of mass of each colony; color indicates mean level of altruism. The two representations overlap in the middle of the figure to demonstrate their consistency. Black squares mark the deaths of colonies; orange circles indicate reproduction of colonies by binary fission. (b) Prediction from linear stability analysis. (See also Appendix B and Fig. S2.) Colonies are expected to emerge in the yellow part of the phase diagram where the scale of competition is clearly larger than the scale of altruism (*σ*_a_ = 1 by definition) and the scale of motility is small. The red cross marks the default parameters used in panel (a). (c) Simulation results testing the prediction of panel (b). As predicted, colonies emerge only in the region to the right of the red line, which is copied from panel (b): the variance of the local population density increases precipitously when the line is crossed. (d) Mean level of altruism at the end of evolutionary simulations. Altruism evolves only in the regime where colonies can form. Each data point plotted represents the mean of three independent replicate simulations. (e) Same as (d), but as a function of mutation probability *μ*. Two independent replicates are plotted in gray; colored circles represent their mean value.

#### Colonies emerge due to a linear instability

The mechanisms behind the emergence of colonies in the 1D habitat can be studied mathematically using linear stability analysis (LSA). We envision a population of individuals with a fixed level of altruism *ϕ* homogeneously distributed over a large habitat that is populated at carrying capacity. Next we superimpose a tiny periodic density variation with some wavelength *λ* and derive under which conditions this perturbation is expected to grow exponentially, resulting in “colonies”. This also allows us to make approximate predictions on the wavelength of the emerging pattern, *i.e*., the distance between colonies. Details are found in Appendix B and Fig. S2.

The LSA reveals that colonies are expected to develop only for certain combinations of the scales of altruism, competition, and motility (Fig. 3c; remember that *σ*_a_ = 1 by definition). As the LSA elegantly demonstrates (see Appendix B for mathematical details) the appearance of colonies is determined by a tug of war between these three forces. Altruism by itself tends to amplify differences in local density: areas with a high density contain more altruists, which positively affects the reproduction rate and hence further increases the density. This ultimately drives the emergence of colonies. However, this force is weak for density variations with a wavelength shorter than ∼ *σ*_a_, which average out within the scale of altruism. Resource competition tends quench density differences because it suppresses reproduction in densely population areas. This force, however, is weak for variations with wavelengths shorter than ∼ *σ*_rc_, which are averaged out within the scale of competition.

Lastly, random motility also tends to homogenize the density, but this force is ineffective against variations with wavelengths larger than ∼ *σ*_m_ because random motion is famously slow at large scales. Together, this means that colonies form only if *σ*_m_ is small compared to the other scales and *σ*_a_ is clearly smaller than *σ*_rc_, so that wavelengths exist that are too long to be quenched by motility, long enough to be amplified by altruism, but too short to be suppressed by resource competition (see Fig. S2a).

To test these predictions, Fig. 3c presents results from a large number of simulations using a range of values of *σ*_m_ and *σ*_rc_ (19 × 21 = 399 simulations in total) in which all individuals are given an immutable value of *ϕ* = 0.05. Each simulation quantified to what extent colonies developed by simply measuring the variance in the local population density. The results are as expected: when crossing over from the linearly stable (blue) to the linearly unstable (yellow) region of Fig. 3b the variance in the local density increases precipitously. The wavelengths of the emerging patterns —typically close to 2*σ*_rc_— also broadly match predictions (Fig. S2c,f). We therefore conclude that the LSA accurately describes and explains the emergence of the colonies.

#### Colony formation by altruists enables evolutionary bootstrapping

From the observations of both the 1D and the 2D model it appeared that the emergence of colonies is important for the evolution of altruism. If so, appreciable levels of altruism should evolve only in the parameter regime where colonies can emerge given reasonable levels of altruism (the linearly unstable, yellow region of Fig. 3b,c). This is confirmed by a series of simulations for various scales of motility and competition (Fig. 3d). Because the colony formation depends on altruism, but the selection of altruism in turn depends on the formation of colonies, the process must pull itself up by the bootstraps. Random mutations plus local reproduction spontaneously result in unstable colonies with modest levels of altruism and high internal levels of drift. This occasionally produces a colony that is altruistic enough to reproduce, which starts to spread rapidly.

Factors other than the spatial length scales clearly also affect whether altruism prevails. If the scale of competition becomes too large relative to the scale of motility, the mean level of altruism suffers (Fig. 3d, *e.g*. at *σ*_rc_ = 5 and *σ*_m_ = 0.1). Also, the stability of colonies against corruption by defectors is affected by the rate with which such defectors are created by mutations. In line with this, the mean level of altruism decreases if the mutation probability is increased (Fig. 3e). That said, altruism emerges for a broad range of mutation probabilities.

#### Colonies die as a consequence of competition

We stressed that even completely selfish individuals with *ϕ* = 0 are viable in isolation. In this light, the observation that colonies characterized by a low mean level of altruism tend to go extinct demands an explanation. The fact that they do implies that the mean reproduction rate of their members sink below the death rate. (Remember that the death rate is independent of environmental conditions.) In the model, only two factors can suppress reproduction: the cost of altruism, and competition. The cost of altruism cannot explain the extinctions because it tends to be low rather than high in colonies with low levels of altruism. By elimination, the extinctions must result from competition with neighboring colonies.

Key to understanding the phenomenon is the observation that colonies with a high level of altruism develop a higher local population density. To good approximation (ignoring motility), the local density settles at a level such that the reproduction rate equals the death rate; if a colony benefits strongly from altruism, this happens at a higher density. As a consequence, a mostly selfish colony that finds itself next to a particularly altruistic one experiences increased competition. This increased competition suppresses its population number. In response, the altruistic colony in fact experience *less* competition, allowing it to expand its population number and to move closer to its neighbor. The latter phenomenon can consistently be seen in Fig. 3a: briefly before a colony goes extinct, neighboring altruistic colonies tend to encroach on it. Thus, sufficiently altruistic colonies can drive sufficiently selfish ones to low population numbers and ultimately extinction.

When a colony is about to disappear, sometimes a few of its surviving members are absorbed by the colony that takes its place. Now infected with defectors, that colony often rapidly declines and ultimately goes extinct too. Under default parameters this process does not allow lineages of defectors to coexist with altruists indefinitely: in simulations in which no new mutations are introduced after the first 80 000 generations, defectors quickly disappear (see Fig. S3).

### 2.4 Quantitative measurement of multilevel selection components

To formally analyze and quantify the role of the colony dynamics in the selection of altruism, we make use of two existing mathematical results. Both are based on subtly different formalizations of the concept of group selection that are sometimes referred to as multilevel selection (MLS) 1 and 2 [13, 41] (see Appendix A for brief derivations).

The two results rely on different applications of the Price equation [40, 42, 43]. The Price equation decomposes the change in the population mean of a trait *ϕ* over a time interval *δ*#x0394;*t*, into two parts: the selection differential *S*, which quantifies the contribution of natural selection, and the transmission term *T*, reflecting systematic differences between the trait value of ancestors and their offspring.

MLS 1 is based on the fact that, in a population that is subdivided into groups, the selection differential *S* can be split into two components: *S* = *S*_within_ + *S*_among_. Here *S*_within_ is the (weighted) mean of the selection differential measured *within* groups, while *S*_among_ is the covariance between a group’s mean trait value and its mean fitness, which can be interpreted as the selection *among* groups.

Our observations suggested that selection within colonies tends to be negative, which is to say that *S*_within_ is systematically negative. To compensate, *S*_among_ would have to be positive as a rule. To test this, we split the simulation illustrated in Fig. 3a into 2000 time intervals of 80 generations each and calculated *S*_within_ and *S*_among_ for each of these time intervals. The results, plotted in Fig. 4a, confirm the expectations. Over the last 1000 time intervals (shaded background in Fig 4), the mean level of altruism no longer changed significantly. (Mean change per time interval is (0.9 ± 2.0) × 10^−5^, where the uncertainty denotes a 95% confidence interval; see Methods.) But over the same period, the within-colony component of selection was negative during 97.6% of the time intervals, averaging (−3.8 ± 0.4) × 10^−4^. In contrast, the among-colony component was positive in 98.7% of the time intervals, with a mean of (4.2 ± 0.4) × 10^−4^. Hence, selection within colonies is indeed negative (reflecting the within-colony tragedy of the commons), but this is compensated by a positive among-colony component of selection.

**Figure 4.**
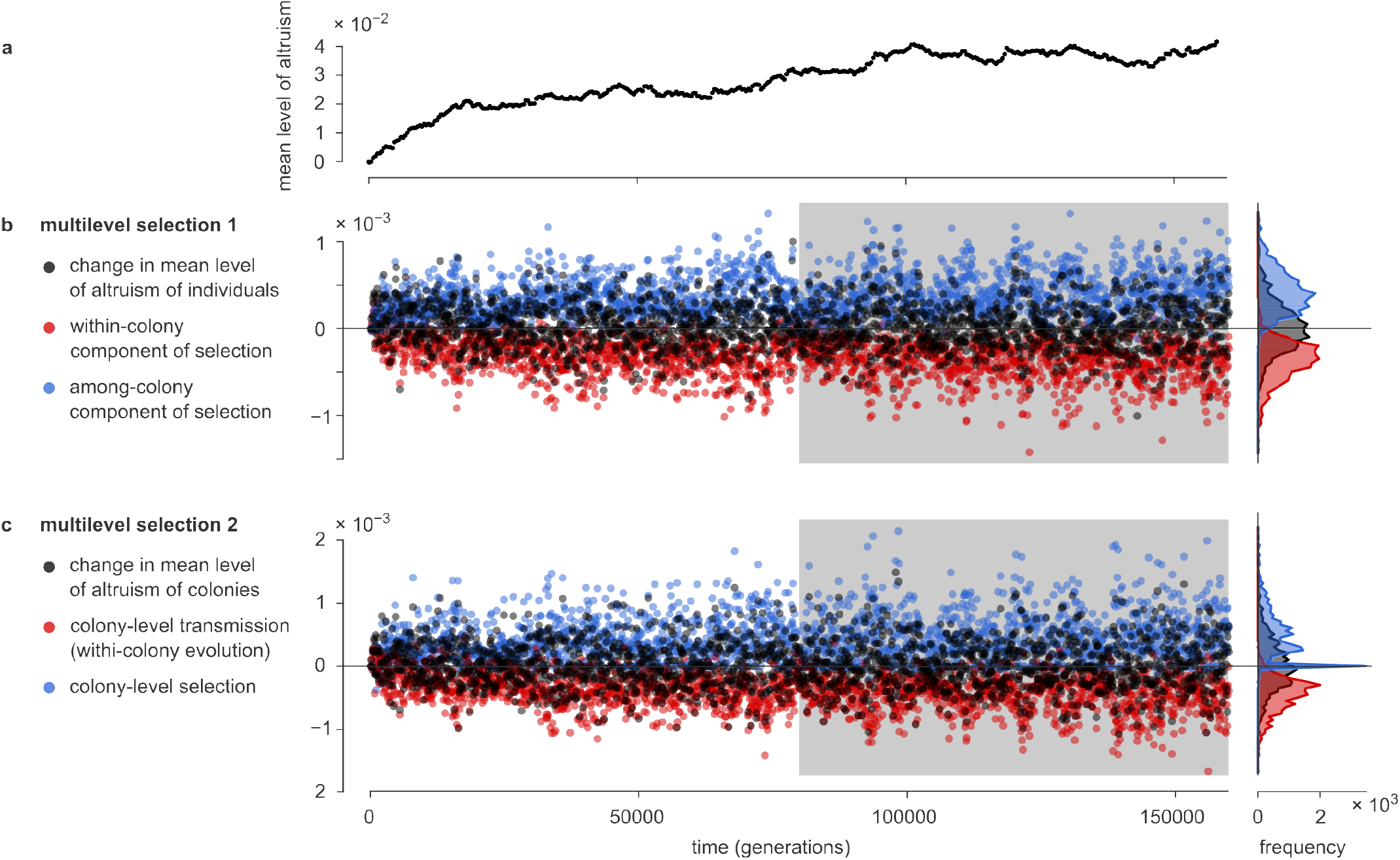
Quantification of multilevel selection: two approaches. Two approaches to quantify multilevel selection are applied to the simulation of Fig. 3a. Fig. S4 shows the results of two more replicate simulations. (a) Population mean level of altruism versus time. (b) The first approach (MLS 1) is to mathematically split the natural selection measured acting on the organisms into two parts: selection within and selection among colonies [40]. (See Appendix A.3.) For the simulation shown in Fig. 3a, this calculation was done for each subsequent interval of 80 generations (see Methods). Plotted is the change in the mean level of altruism (black), and the within-colony (red) and among-colony (blue) components of selection. The rotated histograms on the right-hand side show distributions based on second half of the simulation, indicated with the gray background. In this part of the simulation, the population mean level of altruism no longer changed systematically. However, within-colony selection is nearly always negative, compensated by positive among-colony selection. (c) The second approach (MLS 2) describes evolution at the level of the colonies. (See Appendix A.4.) Evolution taking place within colonies then appears as a transmission bias: a bias in the change between ancestor and offspring colonies in the colony mean level of altruism. This transmission bias (red) tends to be negative, but is compensated by positive selection at the colony level.

The analysis of MLS 1 applies the Price equation to the population of individuals. MLS 2 instead applies it to the population of *colonies*. The mean level of altruism of individuals in a colony, Φ, is now considered a trait of that colony, and the fitness of a colony is defined as the number of offspring *colonies* that it has at the end of the time interval. The Price equation can then be used to describe the change in the mean level of altruism of the colonies. The selection differential S now measures whether colonies with a high value of Φ tend to produce more offspring colonies, and hence can be interpreted as the colony-level selection on Φ. The transmission term T now quantifies to what extent the Φ-value of offspring colonies systematically differs from those of their ancestral colonies. Hence, T characterizes the internal evolution of colonies. (Generally this internal *evolution* cannot directly be interpreted as internal *selection*, as other evolutionary forces may contribute to it.)

It appeared that colonies with a higher mean level of altruism tended to have a higher fitness, suggesting that the colony-level selection is predominantly positive. To test this, we applied the MLS 2 framework to the same 2000 time intervals used for the MLS 1 analysis, making use of the automatically acquired colony-level lineage traces to measure the fitness values of the colonies. The result, plotted in Fig. 4b, again confirms the expectations. Over the last 1000 time intervals, the mean level of altruism of colonies 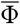 no longer changed significantly. (Mean change per time interval: (0.9 ± 2.0) × 10^−5^.) In the same window, colony-level selection was positive in 82.2% of the time intervals, and negative in only 1.6%. (In the remaining intervals none of the colonies reproduced or died, resulting in a colony-level selection of precisely 0.) Its mean value was (4.7 ± 0.5) × 10^−4^. This was compensated by the colony-level transmission term, which was negative in 97.5% of the intervals, with an average of (−4.6 ± 0.4) × 10^−4^. From this we conclude that the mean level of altruism of individual colonies tends to decrease with time, compensated by an increased fitness of colonies with a higher level of altruism.

We suspected that altruistic colonies are fitter for two reasons: they are *more* likely to reproduce but also *less* likely to die. Mathematically, the relative importance of these two mechanisms can be determined by splitting the colony-level selection of MLS 2 into two terms measuring the contribution of each; see Appendix A.5 for a derivation. We applied this analysis to simulations. As seen in Fig. S5, both terms are overwhelmingly positive, confirming that both mechanisms contribute. That said, the fact that colonies with a higher level of altruism are less likely to die is more consequential by a factor of two.

## 3 Discussion

Above, we have presented a simple model of the evolution of altruism. Despite its simplicity, the model displays complex dynamics. Under suitable parameter settings, a linear instability permits a process of evolutionary bootstrapping in which altruistic colonies emerge that themselves reproduce by binary fission. Quantitative measurements demonstrated that defectors have the upper hand within colonies, but that colonies with a higher mean level of altruism reproduce more frequently and survive longer. The net effect is that a significant level of altruism develops and persists in the population.

Complex biological systems invariably show a hierarchical organization, with collectives of individuals at one level forming entities at a higher level. The evolution of such hierarchical structures involves transitions in which collectives of individuals start to behave as individuals: units of evolution [44] showing group adaptation [45]. An important open question in evolutionary theory is what mechanisms and conditions allow such transitions in individuality to take place [46]. While many theoretical models study evolution in hierarchical population structures, most take this structure as given (e.g., [14, 47]). In some cases, particular parameters of the group structure are allowed to evolve concurrently with the social behavior (*e.g*., [48]), but the group-level life cycle as such is still presupposed. In other models the spatial dynamics spontaneously produce hierarchical structures, but with aggregates that cannot naturally be considered Darwinian entities because they are either too short-lived or do not replicate in a clear-cut sense (e.g., [49–51]). In yet other models, the formation of collectives is initially “scaffolded” by preexisting environmental structure [46]. In this light, a distinguishing feature of the current model is that group-level Darwinian entities emerge spontaneously by self-organization; to our knowledge, few other models have this property (but see [52]). The formation of spatial density patterns is very common in nature and can result from various mechanisms [53]. The model that we presented reconfirms that, even in the absence of a preexisting ecological scaffold, such ecological self-organization can naturally result in competition and replication at the level of aggregates. The aggregates emerging in our model are subject to considerable internal evolution; hence, although most would acknowledge them as units of selection [54], they do not qualify as units of evolution in the sense of Maynard Smith [44, 55] or as a level of adaptation in the sense of Gardner *δ* Grafen [45]. Possibly, however, once natural selection is able to act at the level of such emergent aggregates, this opens an avenue towards a full transition of individuality.

Classical theory argues that the relative scales associated with motility, social interactions and competition are of crucial importance for the evolution of altruism (see Introduction). The results of the model confirm this: as expected, altruism evolves only if motility is limited and the scale of competition is larger than the scale of altruism. The significance of the scales, however, is much more involved than anticipated because they largely determine the emerging ecological patterns, which in turn shape the evolutionary dynamics. Indeed, the evolutionary dynamics support altruism (Fig. 3d) only if the ecological dynamics support the formation of colonies (Fig. 3c). This emphasizes that it is unlikely that the eco-evolutionary behaviors of complex dynamical models can be summarized by generic rules of thumb.

The formation of new colonies through the fission or budding of existing ones is in fact common in social mammals and insects, and its relevance for the evolution of cooperation and altruism has been demonstrated in earlier models (*e.g*., [16, 56–58]). In their formulation, these earlier models tend to differ strongly from the one presented here. They explicitly specify the dynamics of group-level events, such as group extinctions and recolonization by buds or propagules, as well as the effect of social behaviors on these dynamics; in addition, they usually do not explicitly include spatial structure within or among groups. Nevertheless, the emergent dynamics of our model are qualitatively reminiscent of such models. An important lesson drawn from earlier models is that altruistic traits can evolve if they decrease the rate of colony extinctions, increase the recolonization rate of propagules, or permit an increase in colony size [16]. As we have seen, in our model the altruism achieves all three effects to some extent.

Superficially, the model is also reminiscent of the ecological public-good (EPG) games of Wakano *et al* [22], which also produce intricate spatial patterns, including Turing patterns. But upon closer inspection the two models differ fundamentally in multiple aspects. In the EPG model all interactions are entirely local. Its pattern formation depends on a coexistence equilibrium point that has no counterpart in the current model, and parameters are chosen such that defectors are not viable without altruists. In addition, a necessary condition for the Turing patterns of the EPG model is that defectors are more motile than altruists; this distinction does not exist in our model. Importantly, the authors do not report that their colonies replicate, although perhaps such behavior could be obtained in a particular parameter regime. To sum up, the mechanisms producing the spatial patterns qualitatively differ between the two models and the EPG model does not display similar multilevel dynamics.

The concepts of multilevel and group selection and their relation to inclusive fitness theory are the subject of a longstanding and fierce debate [59–63]. Here, we do not engage in this debate. Given the remarkable colony dynamics of the model, a multilevel perspective is particularly natural and allowed us to test relevant hypotheses. But several other theoretical frameworks and fitness-accounting schemes [9] could be applied as well to address different questions. To illustrate this, Fig. S6 presents an inclusive-fitness analysis of one of the 1D simulations with default parameters. The analysis confirms that altruism is individually costly but benefits interaction partners, who tend to be highly related.

In the literature, multiple conceptualizations of multilevel selection exist, including MLS 1, MLS 2, and Contextual Analysis [41, 64]. As to which of these methods best captures the concept of multilevel selection no consensus has yet been reached [13]. Above, we have used the decompositions of MLS 1 and 2 essentially as alternative descriptive statistics, each measuring different but well-defined properties of the system. For example, the colony-level selection term of MLS 2 confirms that more altruistic colonies have more offspring, irrespective of whether one considers this term a proper (or even the sole) measure of the concept of group selection. Similarly, the within-group contribution to selection of MLS 1 is a useful measure to test the hypothesis that selection within colonies is negative on average. We could have applied Contextual Analysis too, to confirm associations between individual or contextual properties and fitness [31, 41]. Their conceptual caveats notwithstanding, each of these methods provides a different vantage point and potentially new insights. The features of our model make it ideally suited to illustrate and test the various approaches to multilevel selection.

Models of evolution in subdivided populations usually assume that the fitness of individuals depends solely on their own traits and those of other group members. Any spatial structure at the sub-colony scale is thus ignored. Moreover, it is typically assumed that groups compete equally among each other, ignoring spatial structure at scales beyond the size of a group. In applications, these assumption are approximations at best, and they certainly do not hold in our model because colonies are not homogeneous and the colony-level dynamics clearly result in spatial assortment of colonies (see Fig. 2b and Fig. 3a). This does not invalidate MLS theory, but serves to remind us that we cannot expect to obtain a complete picture from a single vantage point. In a forthcoming article we describe an complementary multi*scale* approach that allows natural selection to be decomposed into contributions at each spatial scale [65]. This approach can be used to analyze the importance of structures below and beyond the colony level; it is also applicable to models that generate spatial structures such as spirals and waves that are relevant to selection but perhaps too ephemeral to be conceptualized as groups. More generally, over the years many evolutionary concepts have been formalized mathematically [66], but these results are rarely applied to computational and individual-based models. Despite each formalism’s limitations, together they provide a valuable toolbox that allows models to be scrutinized quantitatively from multiple perspectives[41, 67]. We hope that future studies take full advantage of its potential.

## 4 Methods

### 4.1 Detailed description of the model

#### Definition of the model

We envision a population or individuals living in a large habitat, which can be one- or two-dimensional. Each individual is fully characterized by its spatial coordinates plus the value of a single quantitative trait *ϕ*, which indicates its investment in altruistic behavior. The behavior of the individuals is defined by just four stochastic processes: death, motility, reproduction, and heredity with mutation.

**Death** strikes each individual at a fixed (Poisson) rate d; if an individual dies, it disappears from the population. The average lifespan of an individual is *d*^−1^, which we call the *generation time*.

**Motility** is modeled as unbiased diffusion with diffusion constant *k*_D_. It follows that, in a generation time, the root-mean-square displacement of an individual in each spatial dimension is 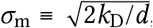, the *scale of motility*. It can be interpreted as the “typical” distance traveled by an individual during its lifetime. Note that we ignore that individuals take up space: nothing prevents multiple individuals from being at the same position at the same time.

**Reproduction** is asexual. When an individual reproduces, a new individual is placed at the same position as the parent. The rate of reproduction of each individual is negatively affected by the level of altruism of the individual itself and by competition for resources with other individuals; in contrast, it is positively affected by the altruism of others in its local environment. To implement these effects mathematically, we make use of two quantities that we will now introduce.

First, we define the local population density *D*(*y*; *σ*_rc_) at position y as a conventional Kernel Density Estimate:

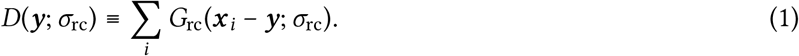

Here, the summation runs over all individuals *i* in the population; ***x***_*i*_ is the position of individual *i*; and the kernel function *G*_rc_(***y***; *σ*_rc_) is the Gaussian (normal) distribution (univariate or bivariate, depending on the dimensionality of the habitat) with standard deviation *σ*_rc_. By this definition, the population density at position ***y*** is high if many individuals are found within a distance of order *σ*_rc_ from ***y***. The parameter *σ*_rc_ is called the *scale of competition* because, as explained below, it determines the range of competitive interactions.

Second, the altruism experienced by an individual at position ***y*** is measured as

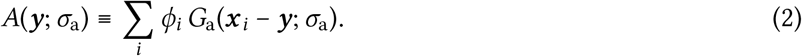

This is again a KDE, except that each individual *i* is weighted by its level of altruism *ϕ*_i_. It is convenient to think of *A*(***y***; *σ*_a_) as the availability of some public good that organisms secrete locally in proportion to their level of altruism. The summation in Eq. 2 runs over all individuals, and the contribution of each individual to the public good at position ***y*** decreases with their distance to ***y*** according to a Gaussian kernel function *G*_a_. The standard deviation *σ*_a_ of the kernel function is referred to as the *scale of altruism* because it determines the range of altruistic interactions.

In terms of these definitions, the full equation for the reproduction rate *g*_*i*_ of individual *i* reads

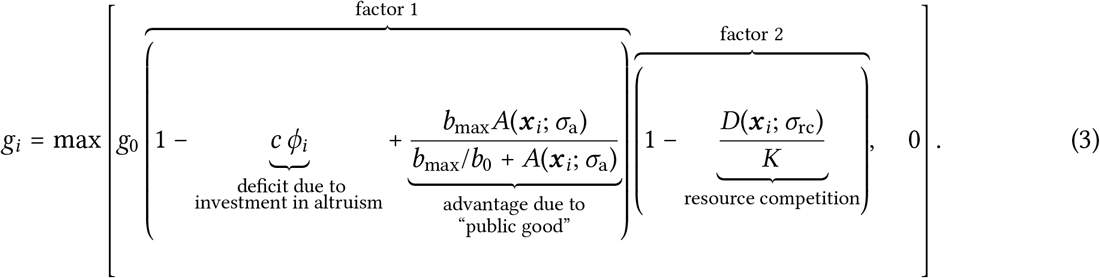

Here, *g*_0_ is the basal reproduction rate. In the subsequent factor (labeled “factor 1”), the term −*cϕ* implements a deficit in the reproduction rate owing to the individual’s investment in altruism; the parameter *c* determines the price of altruism. The last term in factor 1 implements the advantage obtained from the altruism of others. The advantage grows as *b*_0_*A*(***x***; *σ*_a_) if *A*(***x***; *σ*_a_) is small but saturates at *b*_max_ if *A*(***x***; *σ*_a_) is large. Factor 2 introduces resource competition: it decreases linearly with the local population density *D*(***x***; *σ*_rc_) such that reproduction is locally inhibited when the density approaches *K*. (In practice, the population density stabilizes somewhat below *K*, where the average reproductive rate equals the death rate *d*.) The max[.] function is required because both factors 1 and 2 could in rare cases become negative; in that case, *g*_*i*_ is set to 0.

**Heredity and mutation** are implemented as follows. Upon reproduction, the offspring usually inherits the value *ϕ* of the parent, but with probability *μ* a mutation occurs. In that case, the *ϕ*-value of the offspring is determined by adding a random change *δϕ* to the value of the parent. The absolute value of *δϕ* is drawn from an exponential distribution with mean m, and its sign is positive or negative with equal probability. A concern with this procedure, however, is that the resulting trait value *ϕ* of the offspring can become negative. In simulations with a 2D habitat this was not permitted and in such events the value was instead set to 0. Although this is a natural choice, it introduces a mutational bias (see Fig. S1), which would complicate some of the analyses performed on the 1D version of the model (in particular, Fig. 3d,e). In the simulations of the 1D system *ϕ* was therefore allowed to become negative, but the behavior of the individuals was determined by the “effective” value *ϕ*_E_ = max[*ϕ*, 0] rather than by *ϕ* itself. In Fig. 3d,e the mean of *ϕ*_E_ is plotted. In Fig. 3a the colony mean of *ϕ* is plotted, but the distinction is immaterial because in this window of the simulation negative values of *ϕ* are rare.

#### Units and parameter reduction

We are free to choose convenient units for length, time, and the trait *ϕ*; thus, three parameters can be eliminated. First, we choose the unit of length such that the scale of altruism *σ*_a_ equals 1 by definition. The two other length scales that exist in the model, *σ*_rc_ and *σ*_m_, are therefore expressed relative to *σ*_a_. Second, units of time are chosen such that the generation time *d*^−1^ is 1. This implies that the death rate *d* also equals 1 by definition. Third, the unit of the trait value *ϕ* is chosen such that the parameter c (see Eq. 3) equals 1. This simplifies the interpretation of *ϕ*: an individuals with trait value *ϕ* directly sacrifices a fraction *ϕ* of its basal reproductive rate to the public good. Note, however, that the summation in Eq. 2 runs over *all* individuals, so that each individual also benefits from its *own* altruism. In the literature, a distinction is sometimes made between soft and hard altruism, depending on whether the direct benefits that altruists reap from their behavior outweigh the direct costs [47]. We always choose parameters such that the direct costs far exceed the direct benefits, modeling hard altruism. The contribution of individual *i* to the public good *A*(***x***_*i*_; *σ*_a_) at its own position ***x***_*i*_ is given by

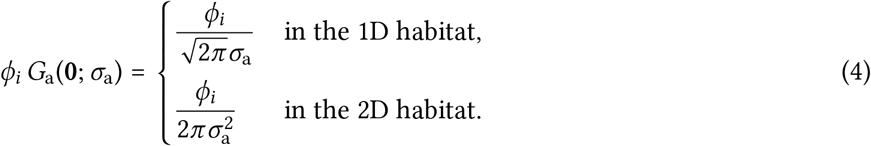

From Eq. 3 and the fact that *σ*_a_ = 1 by definition, it then follows that the reproductive advantage due to one’s own altruism is bounded by 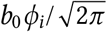 (in the 1D habitat) or *b*_0_ *ϕ* _*i*_/(2*π*) (in the 2D habitat). This reproductive advantage cannot outweigh the deficit of *cϕ*_I_ = *ϕ*_i_ unless 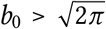 (in the 1D case) or b_0_ > 2*π* (in the 2D case); we steer clear of this regime by choosing *b*_0_ appropriately small.

### 4.2 Implementation of the simulations

#### Simulation scheme

In the simulations, continuous space is approximated by a linear grid (in the 1D habitat) or a square grid (in the 2D habitat) with grid cells of linear size *δ x*. Periodic boundary conditions are imposed. Time is divided into computational time steps *δ t*.

During each computational time step, the state of the system at time *t* + *δ t* is constructed based on the state at time *t* by the following sequence of steps:

**Step 1. Calculate reproduction rates** First, the density *D*(***x*;** *σ*_rc_) and the availability of public good *A*(***x***; *σ*_a_) at each position ***x*** are computed, taking into account the periodic boundary conditions. After this, the reproduction rates *g*_*i*_ of all individuals can be calculated.

**Step 2. Reproduction and mutation** Each individual *i* in the field reproduces with probability *g*_*i*_*δ t*. The offspring is mutated with probability *μ*, as described above.

**Step 3. Death** Each individual subsequently dies with probability *dδ t*.

**Step 4. Motility** Each individual is displaced in each spatial dimension by a distance drawn at random from a discrete approximation of a Gaussian distribution with mean 0 and standard deviation 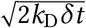.

#### Initial conditions

The steady-state population density of a population of defectors (*ϕ*_*i*_ = 0) is approximately (1 − *d*/*g*_0_)*K*, which can be derived by solving *g*_*i*_ = *d* under the assumption of a homogeneous population distribution. Therefore, the initial condition was constructed by placing (1 − *d*/*g*_0_)*KL* (in the 1D habitat) or (1 − *d*/*g*_0_)*KL*^2^ (in the 2D habitat) defectors at uniformly random positions, where *L* is the linear size of the habitat.

#### Default parameters

The default values of model parameters are listed in Table 1. Here, we also provide the computational parameters used to simulate the model.

For simulations of the 2D model, a square habitat of linear size *L* = 102.4 was used; using *δ x* = 0.1 this amounted to ≈ 10^6^ grid cells. Using the default parameter values *K* = 40 and *g*_0_ = 5, the total population size was approximately n = (1 − *d*/*g*_0_)*KL*^2^ ≈ 3.4 × 10^5^. The simulations were run for *T* = 8 000 generations, with time steps *δ t* = 0.08.

For simulations of the 1D model, a habitat of size *L* = 819.2 was used with *δ x* = 1/80, resulting in 65 536 grid cells. Using the default parameter values *K* = 100 and *g*_0_ = 5, the total population size was approximately (1 − *d*/*g*_0_)*KL* ≈ 6.6 × 10^4^. These simulations were run for *T* = 160 000 generations, again with time steps *δ t* = 0.08.

Additional settings were used to calibrate the automated recognition of colonies; see below.

### 4.3 Computational procedures

#### Calculating cumulative effects of selection, drift, and mutational bias

The mathematical framework used to quantify selection, drift and mutational bias is described in Appendix A.2. We applied this calculation to each time step of the simulations, so that the cumulative effect of each of the evolutionary forces could be tracked (Fig. 2a and S1).

For the analysis we need to obtain, for each individual *i* present right before the computational time step, the expectation value 𝔼 (*W*_*i*_) of the number of offspring *W*_*i*_ it will have after the time step (also counting the individual itself if it survives). To do so, first the growth rate *g*_*i*_ was calculated and subsequently the reproduction and death probabilities *P*_r_ = *g*_*i*_*δ t* and *P*_d_ = *dδ t* over this time step. From the simulation scheme (see above) the expectation value can then be derived:

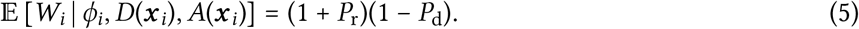

This expression is used in the calculations.

We note that this expectation value is conditioned on the current state of the simulation, in particular the population density *D*(***x***_*i*_) and availability of public good *A*(***x***_*i*_) at the position of the individual. In other words, only the effects of the inherent randomness of reproduction and death given the state of the local neighborhood are accounted as random drift; the fact that the state of the local neighborhood itself is also affected by random events in the past, such as the stochastic motility and demographics of others, is not. (See also Appendix A.2.)

#### Calculating the local population density

To efficiently calculate the local population density (Kernel Density Estimate or KDE) *D*(***x***; *σ*_rc_) (Eq. 1), first a matrix was constructed that specifies, for each position in the habitat, the number of individuals at that position. The KDE at each position, taking into account the periodic boundary conditions, is the circular convolution of this occupancy matrix with the periodic summation of the (discretized approximation of the) Gaussian kernel *G*_rc_(***x***|*σ*_rc_). To perform this convolution, we use the Circular Convolution Theorem, which states that the circular convolution of two matrices can be obtained by first calculating their Discrete Fourier Transform (DFT) and then calculating the inverse DFT of their element-wise product.

#### Calculating the availability of public good

The public good available at each position, *A*(***x***; *σ*_a_), is calculated in a similar way. First a matrix is constructed that contains, for each position in the habitat, the sum of the trait values of all individuals present at that position. The value of *A*(***x***; *σ*_a_) for each position is now obtained as the convolution of this matrix with the (discretized approximation of the) Gaussian kernel *G*_a_(***x***; *σ*_a_), again using the Circular Convolution Theorem.

#### Calculating the radial distribution function

The radial distribution function or pair-correlation function *g*(*r*) is defined as the *observed* number of pairs of individuals separated by a distance *r*, relative to the *expected* number under the null model assuming that each individual is placed at a random position.

In the 1D case, the distance *r* can only take on values *kδx*, where *k* is a non-negative integer. Call the population size *n* and the size of the habitat in grid cells *X*. To calculate the expected number of pairs at distance *r* = *kδx*, written as *E*(*r*), we note that the number of individuals *o*_*x*_ at position x is binomially distributed under the null model. Its expectation value is 𝔼 [*o*_*x*_] = *n*/*X* and hence 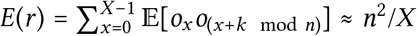. (Here, we used that the occupancies of different sites are to good approximation independent.) The observed number of pairs of individuals found at a distance *r* = *kδx*, called *O*(r), is precisely given by the auto-correlation of the occupancy matrix, which is again efficiently calculated using the Circular Convolution Theorem. For each value *r* = *kδx*, the radial distribution function is then obtained as *g*(*r*) = *O*(*r*)/*E*(*r*).

In the 2D case, the rectangular grid imposes that r can only take on values such that *r*^2^ = (*a*^2^ + *b*^2^)*δx*^2^, where *a* and *b* are integers. In addition, in calculating the expectation under the null model, the frequency *F*(*r*) with which each distance occurs in the grid has to be taken into account. (*E.g*., the distance 5*δx* occurs three times more often than the distance 6*δx*.) Under the same assumptions as made for the 1D case, the expectation is *E*(*r*) = *F*(*r*)*n*^2^/*X*^2^. To calculate the observed number of pairs *O*(r), the auto-correlation matrix of the occupancy matrix is used. Then *g*(*r*) = *O*(*r*)/*E*(*r*) is calculated for each admissible value of *r*. To obtain plot Fig. 2c, the distances were subsequently binned.

#### Calculating the terms of MLS 1 and MLS 2

To obtain Fig 4 and S4, we divided the simulation into 2 000 time intervals of Δ*t* = 80 generations and applied the analyses of MLS 1 and 2 to each time interval. The mathematical expressions for MLS 1 and 2 are briefly summarized in Appendix A.4. Here follows a description of the computational methods used.

For concreteness, let us focus on a particular interval (*t*_1_, *t*_2_]. The first step is to calculate the selection differential *S*, defined as the covariance of *ϕ* and relative fitness *w* (Eq. A.2 in Appendix A.1). For the purpose of this analysis, the relative fitness *w*_i_ of an individual *i* living at time *t*_1_ is the number of offspring it has at time *t*_2_ (the absolute fitness *W*_*i*_), divided by the population mean 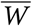. To find these offspring numbers, each individual at time *t*_1_ was assigned a unique ID that was subsequently inherited by all offspring. At time *t*_2_, a frequency table of ID values was constructed, which directly provided the fitness of each each individual at time *t*_1_. With this information, *S* can be calculated directly.

The analysis of MLS 1 splits S into two parts (see Eq. A.8 in Appendix A.3). It is sufficient to calculate the second term, *S*_among_, after which the first follows as *S*_within_ = *S* − *S*_among_. To calculate *S*_among_ first the geographical borders of all colonies at time *t*_1_ were identified; the algorithm used for this is described in the next section. Next, each individual at *t*_1_ was assigned to a colony. Then for each colony j we calculated its population size *n*_*j*_, its mean relative fitness {*w*; *j*}_w_ and its mean trait value {*ϕ*; *j*}_w_. At this point, *S*_among_ could be calculated from its definition.

The analysis of MLS 2 describes the dynamics from the perspective of the colonies (see Eq. A.9). To determine the fitness of the colonies present at time *t*_1_, we had to count how many offspring *colonies* they have at time *t*_2_. This requires that we defined the borders between the colonies at time *t*_2_, but also that we traced the ancestor colony at *t*_1_ for each offspring colony at *t*_2_; the algorithm used is described in the next section. The other ingredient of Eq. A.9 is the trait value Φ of the colonies. Because Φ_*j*_ = {*ϕ*; *j*}_w_ (see section A.4), these quantities were already calculated for MLS 1 and both terms of Eq. A.9 can be evaluated directly.

#### Automated recognition and tracking of colonies

To perform the multilevel selection analysis, the simulation had to automatically recognize colonies and track their ancestry. Because existing clustering algorithms are inefficient for 1D systems and/or difficult to adjust to our needs, we used our own heuristics.

Where to draw the border between neighboring colonies, and when to conclude that one colony has divided into two, is to some degree arbitrary. The results of the analyses, however, do not depend sensitively on such details as long as we use reasonable definitions and apply them consistently.

The basic idea is to identify the borders between colonies with local minima of the population density. However, local minima can also occur temporarily within colonies due to random fluctuations, and such minima should not be confused with true borders between colonies. To solve this, one might exclude local minima if the density at their position exceeds a set threshold, so that only “deep” minima are considered. Such a simple threshold rule can identify most colonies correctly, but issues arise during the binary fission of colonies. During this process the depth of the local minimum that separates the two daughter colonies fluctuates, and hence it is likely to cross the threshold multiple times. Consequently, the threshold rule tends to record multiple events of fission and fusion during a single process of colony division. Similarly, when a dwindling colony is about to disappear, the threshold rule tends to infer series of deaths and resurrections of the same colony.

To prevent this, the algorithm that was used in the simulations in fact uses two density thresholds: a low and a high one, *T*_low_ and *T*_high_. When a new local minimum appears, a new border (and hence the birth of a new colony) is inferred only when the local density at the minimum drops below *T*_low_. In contrast, when an existing border is about to disappear, this is acknowledged only when the density at the associated local minimum rises above *T*_high_. The result is a hysteresis of sorts: when the density of a new minimum drops below *T*_low_ for the first time, a colony is born; if afterwards the density at the border temporarily exceeds *T*_low_, the border is maintained unless it also exceeds *T*_high_.

To be precise, when performing the MLS analysis on the interval (*t*_1_, *t*_2_], the borders of the colonies at time *t*_2_ were constructed as follows:

1. Calculate a smoothed density. The smoothed density at each position was defined as a KDE with bandwidth *σ*_a_/2, taking into account the periodic boundary conditions.
2. Identify local minima. If the smoothed density at grid point *x* is written as *ρ*_*x*_, each x such that *ρ*_*x*_ < *ρ*_*x*+1_ and *ρ*_*x*_ < *ρ*_*x*−1_ marks a local minimum. (Because of the periodic boundary conditions, all indices should be read modulo *X*, the size of the grid.)
3. Determine *tentative* borders between colonies. First, local minima were selected with a density *ρ*_*x*_ < *T*_high_; the other minima were discarded. Each of the selected minimal was then associated with a tentative border which would later be further scrutinized. To ensure that no individuals can sit exactly at a border (causing ambiguity as to which colony it belongs to), borders were positioned *between* grid points. First, the derivative of the density at the position of each minimum was approximated as *ρ*′(*x*) = (*ρ*_*x*+1_ − *ρ*_*x*−1_)/(2*δx*). If a minimum was located at grid point *x*, then a tentative border was placed between x and *x* + 1 if the derivative was negative, and between *x* − 1 and x if the derivative was positive.
4. Assign an ancestor to each tentative colony. The tentative borders implicitly also defined tentative colonies. For each tentative colony at time *t*_2_ an ancestor colony at time *t*_1_ was determined. To do so, we exploited that we have already traced back the ancestry of the *individuals* in the colonies. We then used the expectation that the ancestor colony *P* of a colony *Q* contains most, if not all, ancestors of the *individuals* that belong to *Q*. Based on this, we identified *P* as the ancestor colony that contains the largest fraction of the ancestors of the individuals belonging to *Q*.
5. Reject tentative borders that reflect fluctuations or incomplete divisions. If the colonies on either side of a tentative border had the same ancestral colony, this suggested that a colony division might have taken place. In this case, we compared the density at the corresponding minimum to the low threshold *T*_low_; if the density was above that threshold, the border was rejected. All other tentative borders were now accepted, so that the identification of colonies at *t*_2_ and their ancestor at time *t*_1_ also became final.
6. Count the number of offspring of each ancestral colony. Now that the ancestors of colonies at time *t*_2_ had been identified, the number of offspring—the absolute fitness—of each ancestral colony could be tabulated. If an ancestral colony had fitness 0, it must have died between *t*_1_ and *t*_2_. If an ancestral colony had absolute fitness > 1 it must have divided. (In practice, a fitness above 2 did not occur because Δ*t* is too short to support multiple consecutive divisions.) If a colony had fitness 1, the ancestral colony most likely survived the time interval without reproducing.

The thresholds *T*_high_ and *T*_low_ are parameters; we found that *T*_high_ = 0.7K and *T*_low_ = 0.2K worked well.

#### Inclusive fitness theory and Hamilton’s rule

In Fig. S6 we demonstrate that the simulations can also be analyzed using inclusive fitness theory. To do so, we essentially use a formulation originally due to Queller [68]; we briefly describe it in Appendix A.6. We applied these calculations separately to every computational time step of the simulation.

The method involves a regression variable 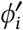, which is usually defined as the sum or the mean of the trait values of all social interaction partners of individual *i*. In our simulation, each individual interact with *all* other individuals, albeit with an interaction strength that decays with their separation (see Eq. 2). We therefore define 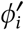 as a weighted sum:

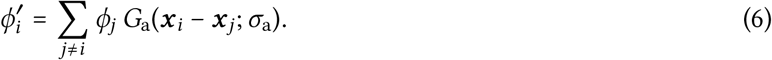

With this definition, the theory of Appendix A.6 can be applied directly.

#### Estimating the lattice constant of the hexagonal lattice by counting colonies

The lattice constant of the hexagonal pattern that emerges in the 2D model can be estimated by counting the number of colonies in the habitat. A hexagonal lattice is composed of equilateral triangles with side *a* and area 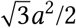. The number of triangles is twice the number of nodes *v*. In a large enough habitat of area *L*^2^, the number of nodes can then be estimated as 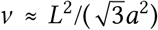. Conversely, after counting the number of nodes, *a* can be estimated as 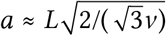. At the end of the simulations of Fig. 2 we find approximately 179 colonies, which, given *L* = 102.4, corresponds to *a* ≈ 8.2. This is consistent with the estimate based on the radial distribution function (Fig. 2c).

#### Estimating error bars

In the Results section and Table S1 we provide 95% confidence intervals for the means of all quantities plotted in Figs. 4 and S4. Because data points in these time series are auto-correlated and the distributions of some quantities are skewed, the standard methods for calculating confidence intervals could not be used. Therefore, we applied the method described in Ref. [69].

Briefly, the idea is to divide the data series into blocks if length *l* and use the means of these blocks (rather than the original data points) to estimate the standard error of the mean (SEM). Starting with *l* = 2, if l is increased, the correlations between block means eventually become negligible and the estimates stabilize around a sensible value, which we determined by manual inspection and then rounded off conservatively. Moreover, because of the central limit theorem, the distribution of block means converges to a normal distribution, which justifies the use of *t*-statistics to estimate confidence intervals. Although the correct number of degrees of freedom to be used is poorly constrained (as it depends on the minimal value of *l* that is deemed large enough to remove correlations), it is in all cases large enough to ensure that the critical *t*-value for t_0.05(2)_ is near 2. We therefore estimated the 95% confidence interval as the sample mean ± twice the estimated SEM.

#### Software

Simulations were performed with custom software written in Fortran; the code is made available on the following GitHub repository: https://github.com/rutgerhermsen/altruism.git. Statistics were performed in R version 3.6.1. Visualization was done in R, using ggplot2, and in Wolfram Mathematica 12.

## 5 Acknowledgments

I am grateful to Rens Dijkhuizen for preliminary simulations and analysis, and to Hilje Doekes for many insightful discussions and valuable feedback. This work was supported by the Human Frontier Science Program, grant nr. RGY0072/2015 (http://www.hfsp.org/funding/research-grants).

## Appendixes

### A The Price equation, evolutionary forces, and MLS 1 & 2

In this article, several mathematical results are applied that have been derived long ago [13, 40, 41]. For ease of reference and to facilitate readers who are not intimately familiar with this theory, we here briefly summarize these results. Nothing in this section is new (with the possible exception of section A.5), although our notation differs somewhat from other presentations to expose the analogies between the multi*level* selection analysis presented here and the multi*scale* analysis presented elsewhere [65].

#### A.1 The Price equation

The Price equation provides a general way to formally describe changes in gene frequencies or mean trait values in evolving populations due to evolutionary forces such as selection and mutation [40, 42, 66].

In its simplest form we consider a population of asexual entities that each possess a numerical trait *ϕ*. At time *t*_1_, the population size is *n*, and the population mean of *ϕ* is 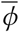. At a later time *t*_2_ = *t*_1_ + Δ*t* the mean of *ϕ* has changed by an amount 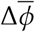. Each individual alive at time *t*_2_ has a unique ancestor at time *t*_1_. (If the individual was already born at time *t*_1_, we designate its past self as the ancestor.) Conversely, each individual i alive at time *t*_1_ has *W*_*i*_ offspring at time *t*_2_. (If the individual is itself still alive at time *t*_2_, it is counted as one of the offspring.) *W*_*i*_ is called the absolute fitness of *i*. The relative fitness *w*_*i*_ of this individual is defined as 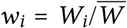, where 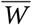 is the population mean absolute fitness. The trait value *ϕ* of the offspring of *i* differs from the value of individual *i* itself; the average difference among *i*’s offspring is called Δ*ϕ*_i_.

With these definitions, the change in the mean value of *ϕ* over the time interval Δ*t* can be written as:

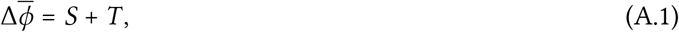

with

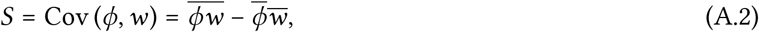

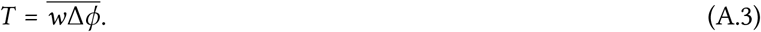

Equation A.1 is called the Price equation. The first term, *S*, is the population covariance between the trait and relative fitness. It shows that the mean value of *ϕ* tends to increase if a high value of *ϕ* is associated with a high fitness. Therefore, *S* is often considered a measure of the effect of natural selection and called the selection differential. The second term, *T*, is the average change in trait value between ancestors and their offspring. Therefore *T* is a measure of transmission bias.

#### A.2 Measuring selection, random drift, and mutational bias

Although the Price equation is frequently and fruitfully used in its standard form, it has its limitations. One clear limitation is that it does not acknowledge one of the evolutionary forces that is central to canonical evolutionary theory: random drift.

The absence of random drift from the standard Price equation is a consequence of the definition of fitness used in its formulation. Above, the fitness *W*_*i*_ of individual *i* was defined as the actual number of offspring it has after the time interval Δ*t*. This is at odds with the usual parlance, in which fitness refers to an organism’s adaptedness to a particular environment. If an organism dies without offspring, this does not necessarily prove that it was poorly adapted to its environment: it might just have been unlucky. The term fitness, then, seems to refer more properly to a propensity or expectation than to an actually realized number of offspring [70, 71]. Deviations from the expectation due to chance are the source of what is usually called random drift.

One way to extend the Price equation is therefore to treat that the number of offspring *W*_*i*_ as a random variable and to associate fitness with its expectation value 𝔼(*W*_*i*_) [13, 66]. In that case we can write the actual number of offspring *W*_*i*_ as 𝔼(*W*_*i*_) + *δ W*_*i*_, where *δ W*_*i*_ is the deviation from the expectation. If we insert this into the standard Price equation (Eq. A.1) we arrive at

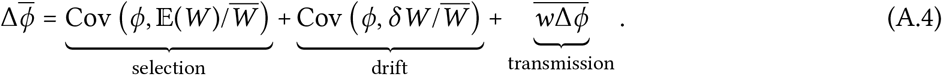

Compared to the standard Price equation, the selection differential *S* is split into two parts: one part that more properly captures the effects of natural selection, and one term that formalizes random drift.

A complication with the above formulation is that it is not obvious how the probability distribution of *W*_*i*_, and hence the expectation 𝔼(*W*_*i*_), should be defined. In particular, it is unclear which variables other than the trait value *ϕ* should be taken into account — that is, which information the probability distribution should be conditioned on. The more information we incorporate into the expectation, the less uncertainty remains to power random drift. Clearly, this difficult issue is beyond the scope of this work. In the meantime, we take a pragmatic stance: Through convenient choices, the above formalism can be used to examine the contributions of elected sources of randomness, regardless of whether these choices can be justified based on unique “correct” definitions of fitness, selection, and random drift.

#### A.3 Multilevel selection 1

Next, we consider a population that is subdivided into *N* distinct groups. To describe the system from the perspective of MLS 1, we start with the Price equation at the level of the individuals, Eq. A.1. The idea of the analysis is to split the selection differential *S* into two parts, *S*_within_ and *S*_among_, where the first accounts for selection taking place *within* groups, and the second for selection *among* groups. We saw that *S* is defined as a covariance (Eq. A.2); mathematically, the decomposition is a direct application of the Law of Total Covariance. In the interest of clarity will nevertheless rederive it from scratch.

It will be useful to introduce some notation. Let *𝓏* be a trait or property of individuals. We will denote the value of *𝓏* of individual *i* in group *j* as *𝓏*_*ij*_, and the size of group *j* will be written *n*_*j*_. Then the mean of *𝓏* within group *j* is written as {*𝓏*; *j*}_w_:

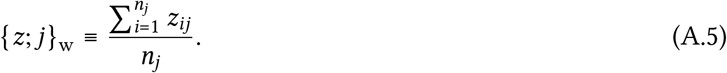

The label “w” stands for “within”. Whenever this does not give rise to confusion we will omit the group index *j* and write {*𝓏*}_w_.

Now, let *u* be a trait or property of groups. Then we define ⟨*u*⟩_a_ as the mean of *u* among groups, where the groups are weighted according to their group size *n*_*j*_:

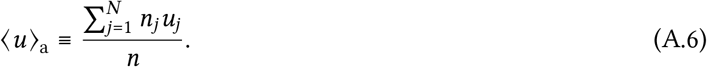

The label “a” stands for “among”.

From the above definitions, one can verify that

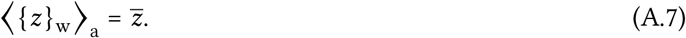

That is to say, if we know the mean value of *𝓏* within each group, {*𝓏*}_w_, we can recover the population mean 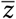 by averaging the over all groups, provided we give larger groups a larger weight.

With the above notation and Eq. A.7 in place, the decomposition of S is obtained quite directly:

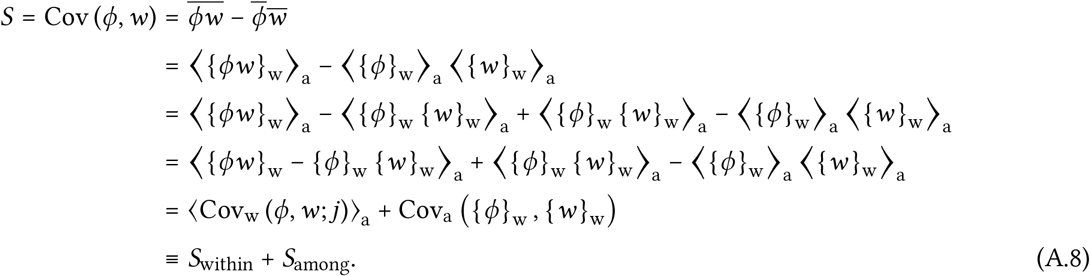

Here we introduced Cov_w_ (*y, 𝓏*; *j*) ≡ {*y𝓏*; *j*}_w_ − {*y*; *j*}_w_ {*𝓏*; *j*}_w_ as the covariance between individual properties *y* and *𝓏* as measured within group *j*, and Cov_a_ (*u, v*) = ⟨*uv*⟩_a_ − ⟨*u*⟩_a_ ⟨*v*⟩_a_ as the covariance of group properties *u* and *v* among groups, where groups are weighted by their group size.

Eq. A.8 shows that *S*_within_ quantifies to what extent within groups the trait value *ϕ* is associated with fitness. It can hence be interpreted as the effect of selection taking place within groups. On the other hand, *S*_among_ measures whether groups with a high mean of *ϕ* tend to have a high mean fitness. It can hence be interpreted as the selection component that results from selection among groups.

#### A.4 Multilevel selection 2

We note that the calculations for MLS 1 can be executed for subdivided populations regardless of whether the groups themselves can in any meaningful way be said to reproduce or die. An alternative formalism, called MLS 2, does explicitly require reproduction at the level of groups.

The idea of MLS 2 is that, if the groups themselves reproduce, the Price equation can be applied at the level of groups. Now the relevant population is the population of groups, and the Price equation can describe the evolution of any trait Φ that is a property of groups:

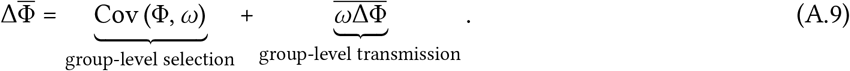

Importantly, the relative fitness *ω*_*j*_ in this Price equation now represents the fitness of group *j*, that is, the (relative) number of groups at time t_2_ that are its offspring (including the group itself, if it survives until *t*_2_).

If we are interested in the evolution of a particular trait at the individual level *ϕ* — such as the level of altruism — we are free to choose Φ to be the group mean of *ϕ*; that is, Φ_*j*_ = {*ϕ*; *j*}_w_. The first term in Eq. A.9 then measures the effect of selection at the group level on the mean trait value of groups. The second term quantifies the effect of bias in the changes in Φ between ancestral groups and their offspring; this reflects the internal evolution of groups.

#### A.5 Separating contributions to selection of reproduction and death

We saw that the Price equations is always applied to a particular time interval (*t*_1_, *t*_2_]. The fitness of an organism or group at time *t*_1_ was defined as the number of offspring it has at time *t*_2_. These fitnesses are therefore determined by two types of events: reproduction and death. It seems clear that natural selection could result from the association of a trait with either of these types of events; hence, we should be able to separate the effects of death and reproduction on selection.

To do so, we note that absolute fitness *W*_*i*_ of individual *i* can be written as follows:

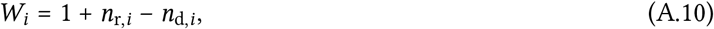

where *n*_r,*i*_ and *n*_d,*i*_ respectively represent the number of reproduction and deaths events occurring withing the lineage starting with individual *i* during the time interval (*t*_1_, *t*_2_]. (Here, events involving i itself should be counted as well.) Inserting this into the expression for the selection differential 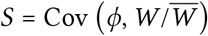 directly gives:

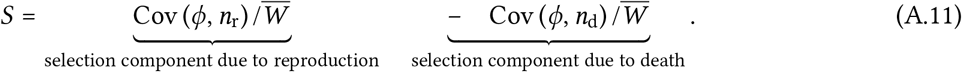

These two terms measure whether the trait is associated with reproduction or death, respectively.

Eq. A.11 dissects selection at the individual level, but if we substitute individual-level quantities by their group-level analogues, the same analysis can be applied to the group-level selection term of MLS 2 to assess the effects of death and reproduction of *groups* on the selection of *groups*.

#### A.6 Inclusive fitness theory and Hamilton’s rule

Within inclusive fitness theory, a variety of approaches and formalisms have been developed [1, 2]. Here we will merely derive the approach used for the analysis of Fig. S6, which is based on Ref. [68] by Queller.

The idea is to write relative (neighbor-modulated) fitness in the following linear-regression form:

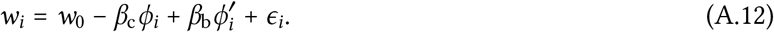

Here, *ϕ*_*i*_ is the trait value of individual i, and 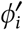 is the sum of the trait values of all individuals that *i* interacts with. The intercept *W*_0_ and the partial regression coefficients *β*_*i*_ are obtained by minimizing the sum of the squared residuals *ϵ*_*i*_.

It is important to note that we do *not* assume that that *w* truly is a linear function of *ϕ* and *ϕ*′. Whether or not w is linear, the linear fit of Eq. A.12 always exists (provided *ϕ* and *ϕ*′ vary within the population and are not collinear); non-linearities are absorbed by the residuals. Also, we do *not* assume that the residuals are independent and normally distributed. Such assumptions are often made in the context of statistical hypothesis tests based on linear models because they conveniently ensure that the desired *p*-values can be calculated from *t* or *F* distributions. Here, it is not our intention to perform a statistical hypothesis test and hence there is no reason to make these assumptions.

With this in mind, parameter *β*_c_ expressed whether, as a trend, fitness tends to increase or decrease with *ϕ* if *ϕ*′ is kept constant. It can therefore be interpreted as a measure of the cost of the trait. (The minus sign in Eq. A.12 was chosen to ensure that a positive *β*_*c*_ indicates that the trait comes at a cost.) Similarly, *β*_b_ measures to what extend fitness tends to increase with *ϕ*′ (as *ϕ* is held constant) and hence summarizes the benefit of the trait to social partners.

Next, we insert Eq. A.12 in Eq. A.2, the expression for the selection differential S. The term Cov (*ϕ, ϵ*) vanishes because, as a general property of least-square linear fits, the covariance between the residuals and each of the regression variables is zero. The result is:

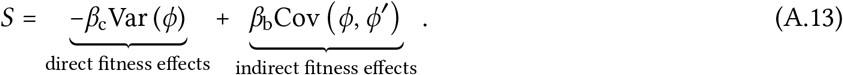

The first term in Eq. A.13 represents the component of selection resulting from direct fitness effects, *i.e*., costs to the actor. The second term represents selection resulting from indirect fitness effects.

A trait is under positive selection if *S* > 0. From Eq. A.13 this can be rewritten as

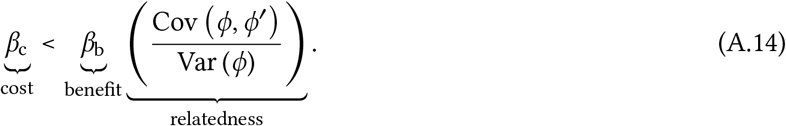

The factor labeled “relatedness” takes the form of a regression coefficient between the trait value *ϕ* of the actor and the total trait value of its interaction partners *ϕ*′. Hence, this is a version of Hamilton’s rule. It states that a costly social trait can nevertheless be selected provided the benefits are large enough and accrue disproportionately to individuals with a high trait value.

If individuals interact with multiple interaction partners, *ϕ*′ can on average be considerably larger than *ϕ*, so that the relatedness factor in Eq. A.14 may become (much) larger than one. In addition, it can in theory assume negative values. This is in contrast to some popular alternative versions of Hamilton’s rule, in which relatedness is confined to the interval [0, 1].

### B Linear stability analysis

We here provide the details of the linear stability analysis for the 1D habitat that is presented in Fig. 3b,c and Fig. S2.

#### B.1 Mathematical analysis

Consider a population of individuals that each have the same level of altruism *ϕ*. If the carrying capacity is large, the dynamics if the density *ρ*(*x, t*) can be approximated by the following mean-field equation:

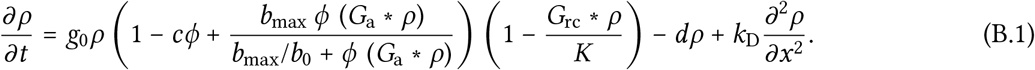

Here, *G*_a_ and *G*_rc_ are the kernel functions used in Eq. 1 and Eq. 2 to define the availability of public good and the local density, respectively. The notation *f* ∗ *h* stands for the convolution of functions *f* and h. Eq. B.1 has a homogeneous equilibrium solution *ρ*(*x, t*) = *ρ*_0_ > 0; we ask under what conditions this solution is (linearly) unstable to periodic perturbations so that colonies can form spontaneously.

To find out, we first identify *ρ*_0_ by equating Eq. B.1 to zero and solving for *ρ* (*x, t*) = *ρ*_0_. Ignoring the trivial solution *ρ*_0_ = 0, the equation is quadratic and can be solved straightforwardly. Out of the two solutions, one is negative and hence irrelevant. The remaining solution depends on all parameters except for the diffusion constant *k*_D_.

We then consider a periodic perturbation

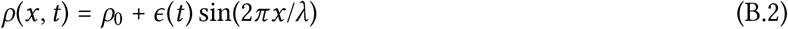

with a very small (infinitesimal) amplitude *ϵ*(0) and ask whether *ϵ* (*t*) will grow or decay. To obtain a dynamic equation for *ϵ* (*t*) we insert Eq. B.2 into Eq. B.1. In doing so, we have to work out the convolutions of the Gaussian kernel functions *G*_a_ and *G*_rc_ with the sine wave of Eq. B.2. From the Convolution Theorem it follows that, for any real-valued, normalized, symmetric kernel function *f* (*x*) that has a Fourier transform f 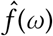, the convolution with a sine wave is again a sine wave, but with a reduced amplitude:

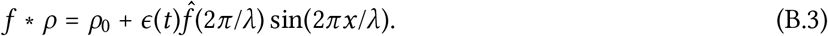

In the specific case where *f* (*x*) is Gaussian with standard deviation *σ*, we get

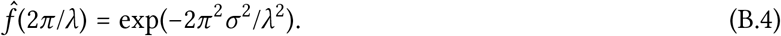

The convolutions with *G*_a_ and *G*_rc_ follow directly from Eq. B.3 and Eq. B.4.

We then expand the resulting equation to first order in *ϵ*(*t*). Because *ρ*_0_ is the homogeneous solution, the zeroth-order term vanishes. The result is a linear equation of the form:

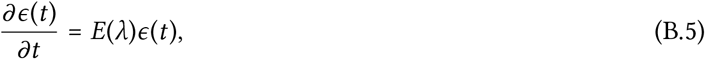

where the factor *E*(*λ*) can be written as:

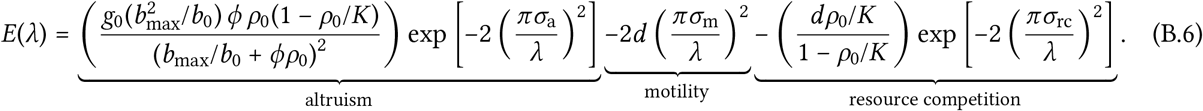

The solution of Eq. B.5 is exponential, with growth rate or eigenvalue E(*λ*). Hence, if Eq. B.6 is positive for some wavelength *λ*, perturbations with this wavelength are predicted to grow exponentially. Because demographic noise produces perturbations of any wavelength, this is expected to eventually give rise to periodic density fluctuations with a similar wavelength.

Eq. B.6 provides considerable insight. It consists of three terms, expressing the effects of altruism, motility, and resource competition, as indicated. The three length scales in the system —the scale of altruism *σ*_a_, the scale of motility *σ*_m_, and the scale of competition *σ*_rc_— each appear in their appropriate term.

The first term, describing the effect of altruism, is the only positive one, and it scales with *ϕ*. This shows that altruism is required to obtain a positive eigenvalue for any wavelength *λ*. Indeed, altruism tends to amplify density differences: because the benefits of altruism grow with the number of altruists in the local neighborhood, its effect is to increase the reproduction rate in regions of high density, which tends to further increase that density. However, the equation shows that this positive contribution is exponentially suppressed if the wavelength *ϵ* is small relative to the scale of altruism *σ*_a_; this is because the kernel function averages out such short waves within the social neighborhoods of individuals.

The second term reflects the effect of motility. Random motion (diffusion) is a homogenizing force and therefore quenches density fluctuations, as reflected in the negative sign of this term. However, because diffusion is famously slow on large length scales, only short wavelengths are strongly affected: if the wavelength *λ* exceeds the scale of motility (the typical distance traveled by an individual in a generation time) the contribution becomes small.

The third term describes the effect of resource competition. Resource competition reduces the reproduction rate in areas with a larger density and thus suppresses density differences, which explains that its contribution is negative. However, if the wavelength is small relative to the scale of competition *σ*_rc_, the density wave averages out within the competitive neighborhood of individuals and the homogenizing effect becomes weak.

Together, this clearly indicates in which regime we ought to expect colonies. Density fluctuations are suppressed by diffusion if their wavelength *λ* is smaller than *σ*_m_, and by resource competition if *λ* is increased far beyond *σ*_rc_. Instabilities are therefore expected only if there is a gap between these two regimes. At the same time, the positive contribution of altruism becomes weak if *λ* is smaller than *σ*_a_. For altruism to be effective in the “gap”, *σ*_a_ therefore should be chosen smaller than *σ*_rc_. In summary, instability requires that the diffusion constant is small enough, the scale of competition is large enough, and the scale of altruism is smaller than the scale of competition. Apart from these rules of thumb, Eq. B.6 can of course be evaluated numerically to make precise predictions; see Fig. S2.

#### B.2 Validation of predictions

In Fig. 3b,c and Fig. S2 we test predictions based on the linear stability analysis using simulations. The key predictions are (i) the region of parameter space where colonies can form, and (ii) the wavelength of the resulting pattern. The following methods were used.

As illustrated with the red dot and arrow in Fig. S2a, both predictions are found by maximizing *E*(*λ*). Fig. S2b shows a contour plot of the maximal value of *E*(*λ*) under variation of the scales; Fig. S2c the corresponding wavelengths. Both values were obtained by differentiating Eq. B.6 and numerical root finding.

To test the predictions we performed a large number of simulations using different values for *σ*_rc_ and *σ*_m_. (We used *σ*_rc_ ∈ {1, 1.2, 1.4, …, 5} and *σ*_m_ ∈ {0.0671, 0.100, 0.134, …, 0.671} in all 21 × 19 = 399 combinations.) As a simple proxy for the presence of colonies, we measured the variance of the (smoothed) population density (KDE) over space. To identify the dominant wavelength in the density pattern, we calculated the Fourier transform of the KDE and selected the mode with the largest amplitude.

The simulations were performed as usual and using default parameters except for the following adjustments:

1. All individuals were initialized with a trait value *ϕ* = 0.05.
2. In these simulations, we were interested in the ecological patterns of a population with fixed *ϕ*; therefore the mutation rate was set to *μ* = 0 to disable evolution.
3. The simulation was run for *T* = 2 400 generations. (The colonies establish very rapidly.)
4. Starting at *t* = 400 generations, after each time interval of 80 generations the following analysis was performed:
  a. Calculate a KDE using a Gaussian kernel with standard deviation/bandwidth *σ*_a_/2.
  b. Calculate the variance of this KDE.
  c. Calculate the Fourier transform of the KDE and identify the wave number with the largest amplitude.

After the simulation, the mean value of the variance was reported. It is this variance that is plotted in Fig 3c and Fig. S2e. Also, the mean value of the wave number with the largest amplitude was reported; this wave number was transformed to a wave length, which was plotted in Fig. S2f.

## Supplementary Figures

**Figure S1.**
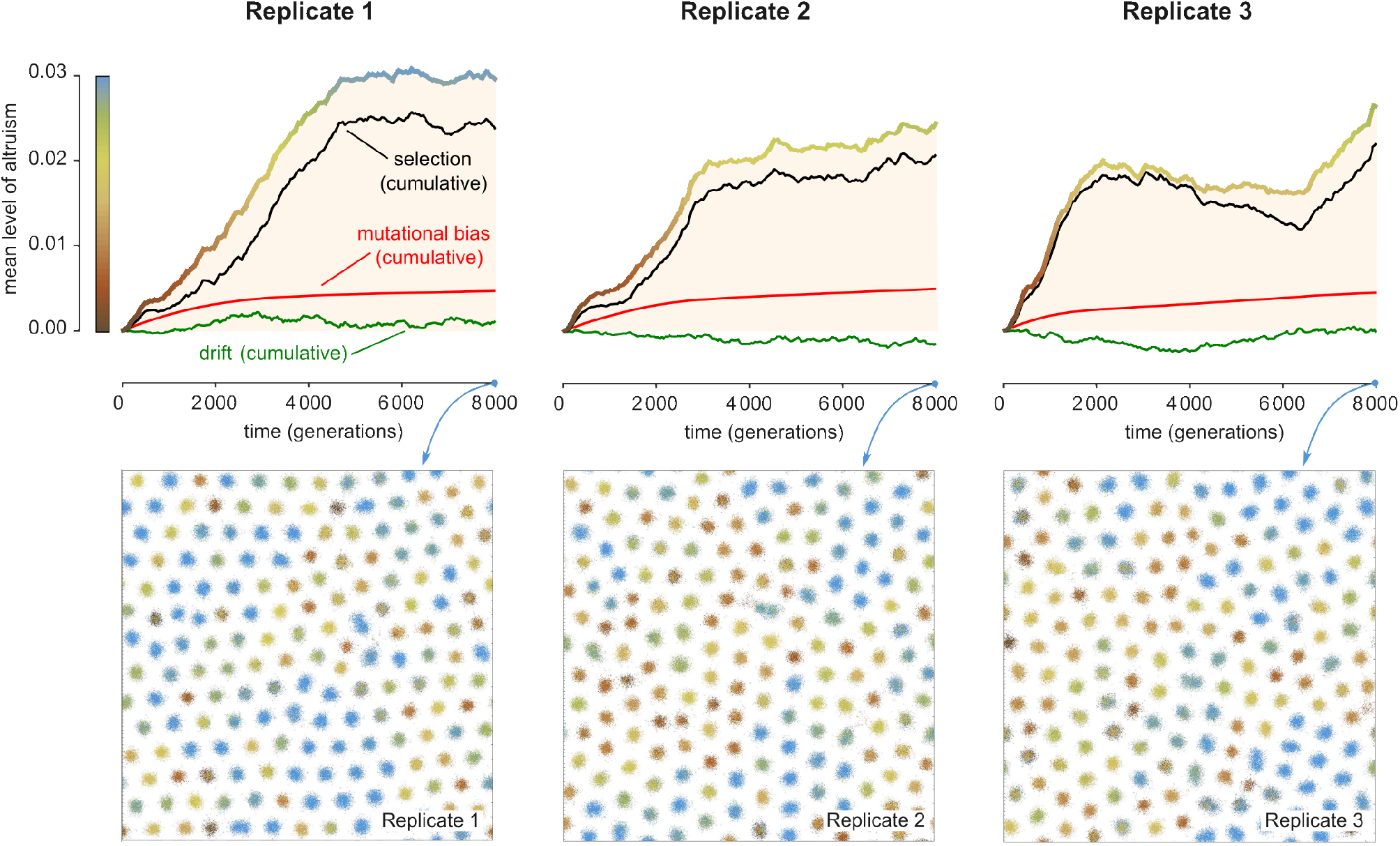
Evolutionary forces and snapshots of replicates (2D habitat) Results are shown based on three simulations that were identical except that the random-number generator was initiated with different random seeds. Fig. 2 presents results of Replicate 1. The three figures on top show the mean level of altruism through time (thick colored line). The rise in mean level of altruism can be decomposed into three contributions: natural selection, mutational bias, and random drift, using the method explained in Appendix A.2. Plotted are the cumulative contributions of natural selection (black) transmission (red) and genetic drift (green). In all cases, the main contribution is selection, which is consistently positive during the first stretch of the simulations. That said, a mutational bias is revealed as well (red lines). This bias arises because, in this simulation, negative values of *ϕ* were prohibited (see Methods) and hence mutations with negative effect are sometimes truncated, especially in individuals with a low trait value. (The smoothness of the red line is a result of the law of large numbers.) The cumulative effect of random drift (green line) is minor in all three replicates. The three figures at the bottom show snapshots of the population at the end of the simulations. In all three cases a hexagonal pattern of colonies has emerged.

**Figure S2.**
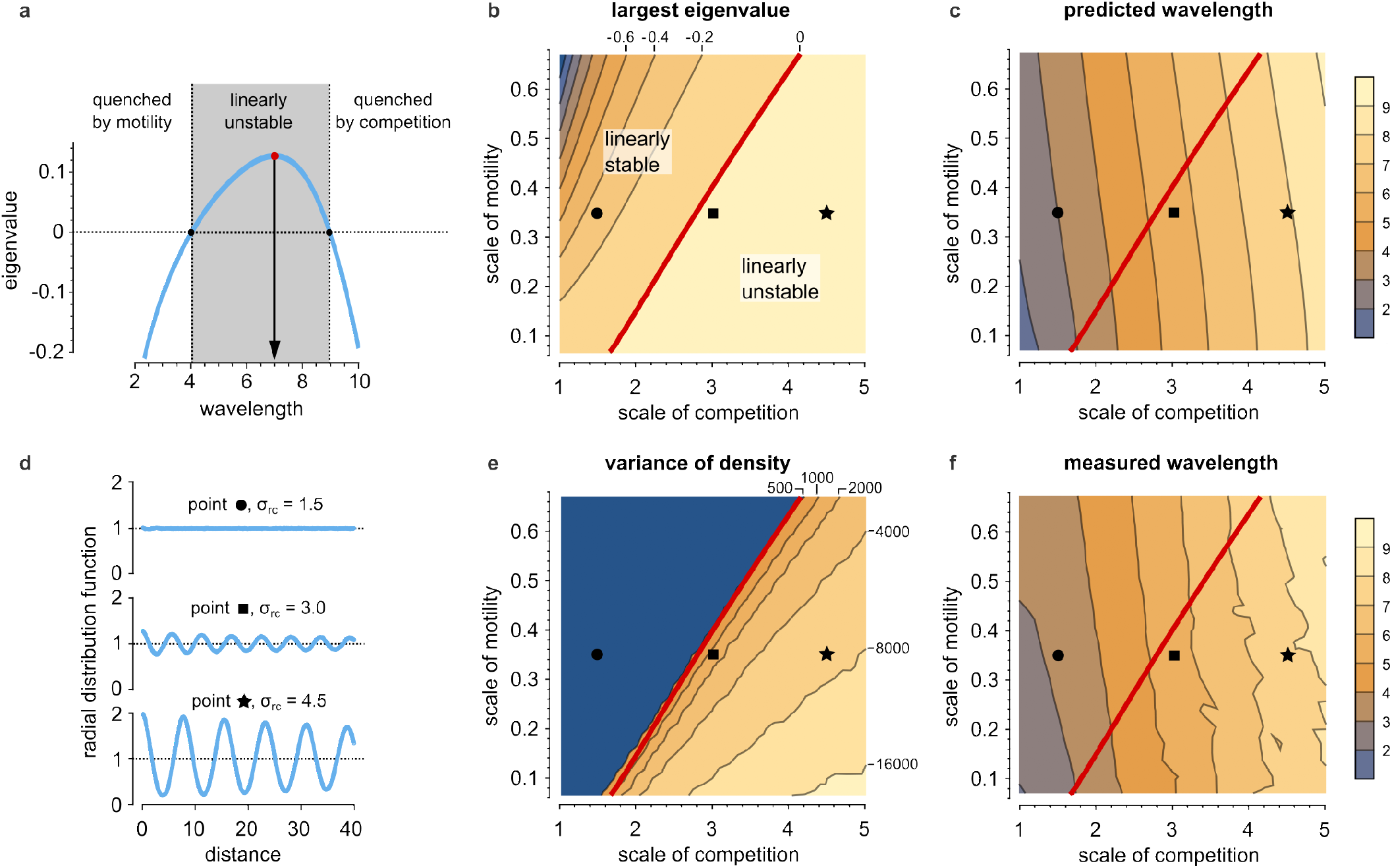
Linear stability analysis. To help understand the conditions for the formation of colonies, a linear stability analysis studies whether, in an initially homogeneous population, small periodic density perturbations tend to grow. (See Methods for the full derivations.) If they do, this leads to the formation of “colonies”. (a) For given model parameters, each wavelength is associated with an eigenvalue; if the eigenvalue is positive, density waves with this wavelength tend to grow. The figure plots the eigenvalue for a range of wavelengths as calculated for the default parameters of the 1D model (Table 1), additionally assuming all individuals have *ϕ* = 0.05. Perturbations with small wavelength are quenched by motility; those with a long wavelength by resource competition. In between, a window exists (gray shading) of wavelengths that have a positive eigenvalue. This explains the colony formation in the default parameters. The wavelength with the largest eigenvalue (indicated with the red dot and black arrow) provides a prediction for the wavelength —the distance between neighboring colonies. (b) The largest eigenvalue is plotted as a function of the spatial scales in the system: the scale of motility *σ*_m_ and the scale of competition *σ*_rc_. (Remember that the scale of altruism *σ*_a_ is 1 by definition of the unit of length.) Otherwise, the assumptions are as in panel (a). Colony formation is expected only in the linearly unstable regime, to the right of the red contour line. (c) For the same conditions used in panel (b), the predicted wavelength is plotted. As a rule of thumb, it is somewhat smaller than 2*σ*_rc_. (d) Simulations were performed for the parameters indicated with black symbols in panels (b), (c), (e), and (f), assuming that all individuals have an immutable level of altruism *ϕ* = 0.05. Shown are the resulting radial distribution functions. As expected, the system is near homogeneous at *σ*_rc_ = 1.5 (black circle, in the linearly stable regime), weak pair correlations are seen for *σ*_rc_ = 3.0 (marginally unstable), and strong correlations emerge for *σ*_rc_ = 4.5 (far in the unstable regime), indicating colony formation. (e) Simulation were performed for a large number of combinations of the spatial scales (19 × 21 = 399 in total); here, the variance of the local population density is plotted as a simple proxy for colony formation. Clearly, colonies form only in the parameter regime predicted in panel (b). (f) For the simulations of panel (e), the wavelength of the resulting colonies is plotted. (See Methods.). The results agree broadly with the predictions of panel (c).

**Figure S3.**
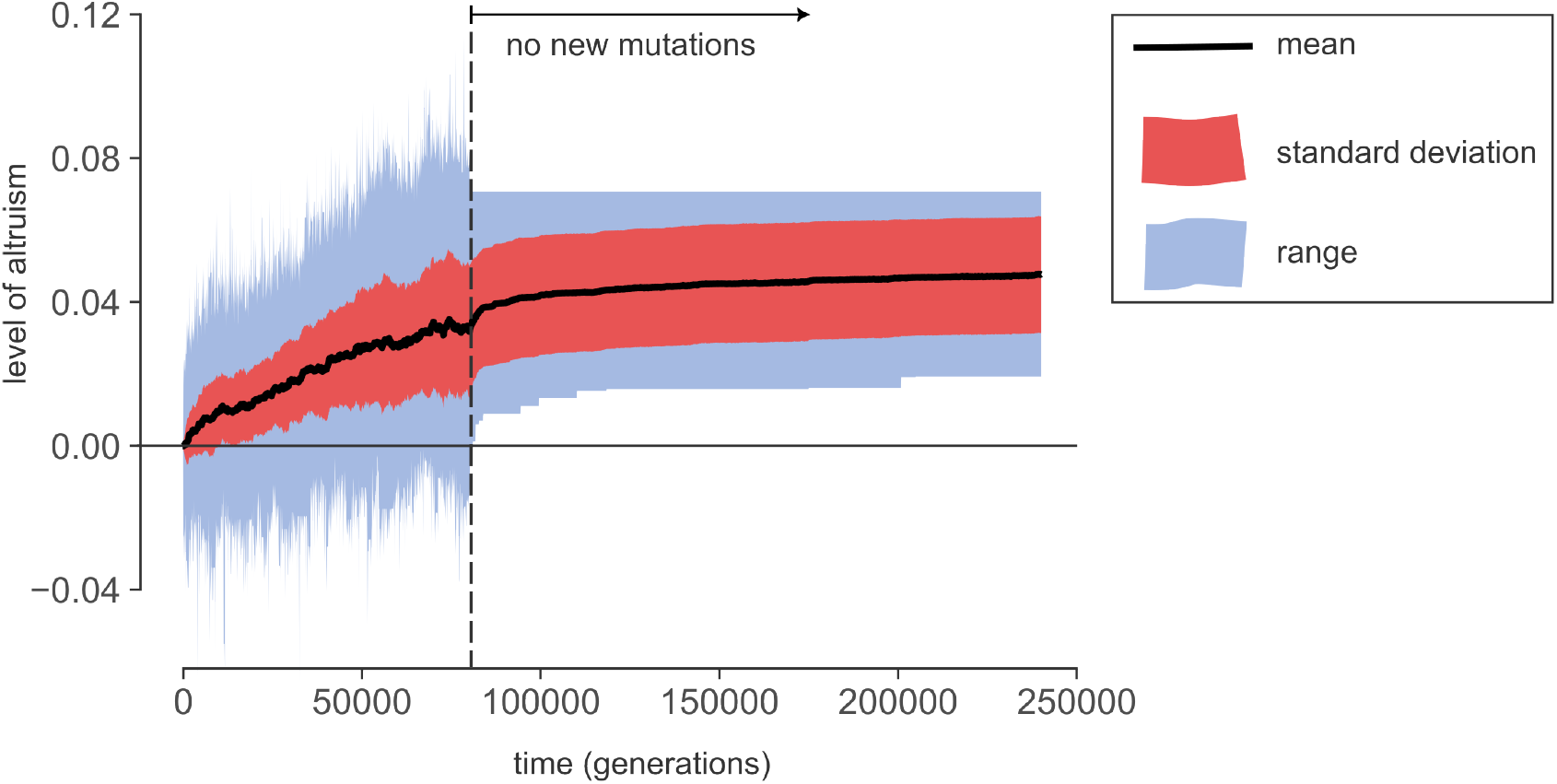
Mutations are required to maintain defectors. Results are shown for a representative simulation of the 1D model with default parameters. In contrast to other simulation runs, no new mutations are introduced after *t*_nm_ = 80 000 generations (vertical line). As a result, directly after *t*_nm_ the range of trait values (blue band) rapidly contracts and defectors (with level of altruism below ∼ 0.01) are eliminated. After this initial decline the distribution of trait values hardly changes: without mutations, the colonies become highly stable and hence between-colony variability is maintained.

**Figure S4.**
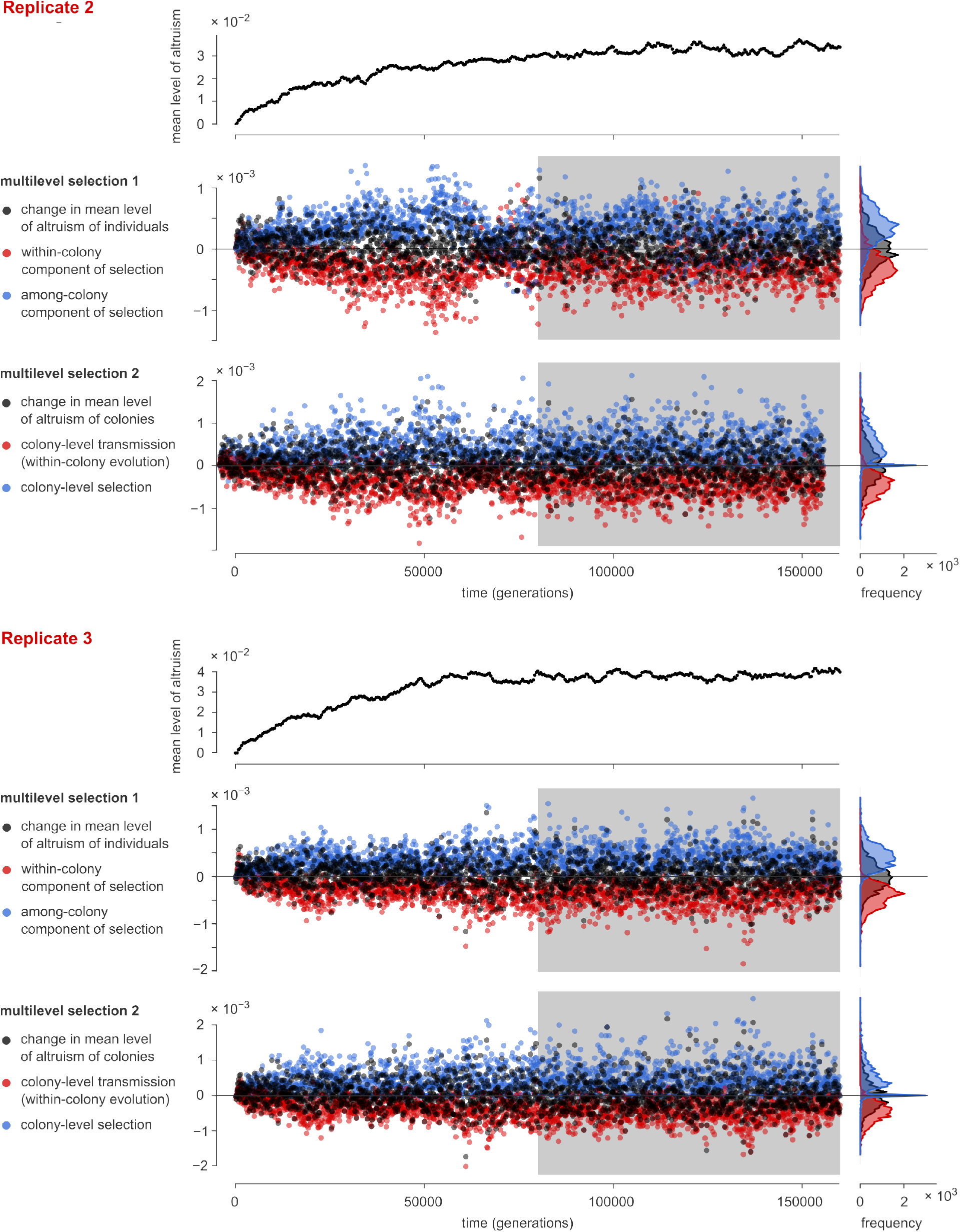
Quantification of multilevel selection (MLS) in replicates (1D habitat). Fig. 4 shows results of the quantification of MLS in a single simulation run. To demonstrate the reproducibility of these results, this figure shows the same analysis for two additional replicates. The simulations for all three replicates were identical except that the random-number generator was initiated with a different random seed. All replicates show very similar trends. In particular, the marginal distributions of all quantities are highly consistent. Their statistics are summarized in Table S1.

**Figure S5.**
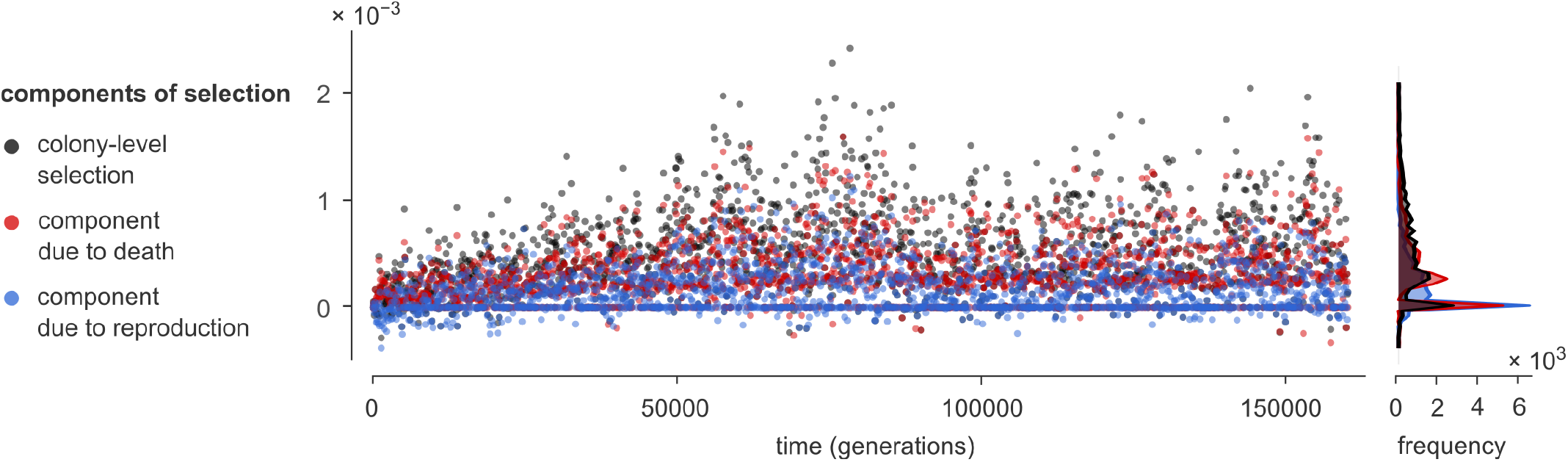
Separating the effects of reproduction and death to colony-level selection. Two mechanisms contribute to the selection of altruistic colonies: they reproduce more frequently and die less frequently. To evaluate the relative importance of these mechanisms we quantify their contribution to the colony-level selection as defined by the MLS 2 analysis; see Appendices A.4 and A.5 for mathematical details. Shown are the results of a single representative simulation with default parameters. As in Fig. 4, the analysis was applied to subsequent time intervals of 80 generations (1 000 computational time steps); the figure plots the total colony-level selection (black) as well as the components due to death (red) and reproduction (blue) events. The marginal histograms on the right-hand side are based on the second half of the simulation, after generation 80 000. Note that the components can be precisely zero if no death and/or reproduction events occur in the particular time interval, resulting in considerable overplotting. First, the results demonstrate that both components to selection are much more likely to be positive than negative (Binomial test, *p* < 10^−15^ in both cases). The component due to death events was positive in 72.6% and negative in just 1.0% of the time steps, with an average value of (3.2 ± 0.4) × 10^−4^. The component due to reproduction events was positive in 62.3% and negative in 8.5% of the time steps, with an average value of (1.6 ± 0.2) × 10^−4^. We conclude that both mechanisms contribute to colony-level selection, but the component due to death is the larger by a factor of two.

**Figure S6.**
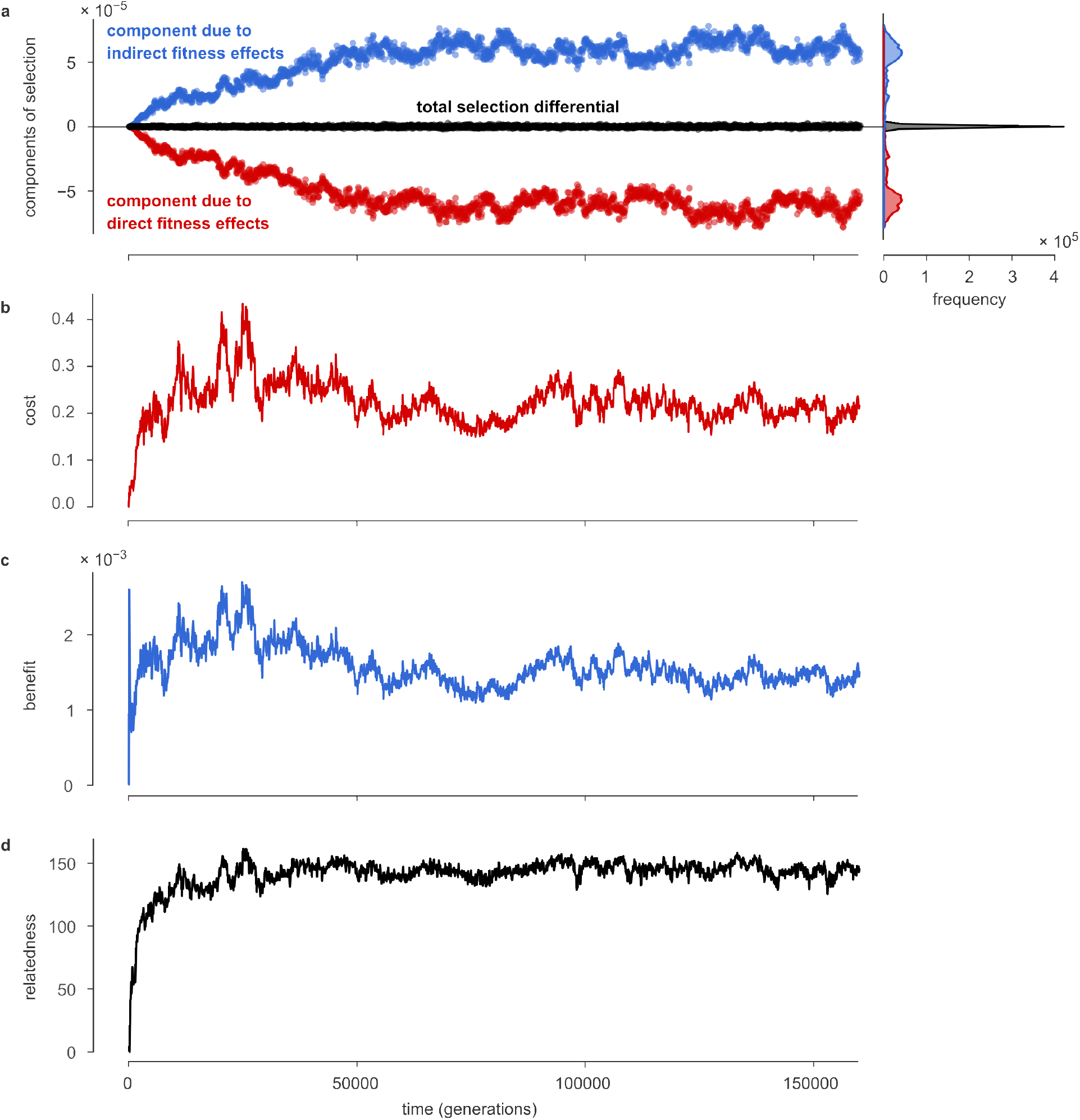
Application of inclusive fitness theory. Figs. 4 and S4 presented an analysis of the simulations of the 1D model based on multilevel selection theory. Here, it is demonstrated that such simulations can also be analyzed from the perspective of inclusive fitness theory. There are multiple ways to do so; here we essentially adopt a formalism based on a partial regression analysis [68]; see Methods and Appendix A.6 for details. Shown are the results of a single representative simulation with default parameters. In all panels, each data point represents an average of the plotted quantity over the preceding 1000 computational time steps (corresponding to 80 generations). (a) The method splits the selection differential S into two parts. The first part, plotted in red, reflects the selection due to the direct fitness effects: the cost of altruism for the actor. This term is consistently negative, which means that altruists face negative selection due to the direct fitness effects, as expected. The second part, plotted in blue, reflects the selection component due to indirect fitness effects: the benefits of altruism for the recipient. That this term is consistently positive indicates that the benefits must preferentially accrue to more altruistic individuals. Both terms approximately cancel out: the sum of both components *S* (black) is positive on average but much smaller in absolute value than either term separately. (b) The cost appearing in Hamilton’s rule, measured as a (partial) regression coefficient (Appendix A.6), as a function of time. This cost is consistently positive, confirming that altruism is individually costly. (c) The benefit appearing in Hamilton’s rule, demonstrating (as expected) that interacting with altruists yields a benefit. (d) Relatedness among interaction partners is consistently positive. We note that, in the formulation of inclusive fitness theory used here, relatedness can legitimately become much larger than 1 (see Appendix A.6).

## Supplementary Tables

**Table S1.** Statistics of the multilevel selection analysis of Figs. 4 and S4. Error bars for the means are given as 95% confidence intervals (see Methods).

| MLS 1 | quantity | replicate | % positive | % negative | mean | 95% CI |
| --- | --- | --- | --- | --- | --- | --- |
| | change in mean of $\phi$ | 1 | 49.4 | 50.6 | $0.9 \times 10^{-5}$ | $\pm 2.0 \times 10^{-5}$ |
| | | 2 | 49.4 | 50.6 | $0.4 \times 10^{-5}$ | $\pm 1.8 \times 10^{-5}$ |
| | | 3 | 48.2 | 51.8 | $-9.1 \times 10^{-7}$ | $\pm 1.7 \times 10^{-5}$ |
| | within-colony component of selection | 1 | 2.4 | 97.6 | $-3.8 \times 10^{-4}$ | $\pm 0.4 \times 10^{-4}$ |
| | | 2 | 10.8 | 89.2 | $-3.2 \times 10^{-4}$ | $\pm 0.4 \times 10^{-4}$ |
| | | 3 | 2.2 | 98.8 | $-4.1 \times 10^{-4}$ | $\pm 0.4 \times 10^{-4}$ |
| | among-colony component of selection | 1 | 98.7 | 1.3 | $\pm 4.2 \times 10^{-4}$ | $\pm 0.4 \times 10^{-4}$ |
| | | 2 | 90.1 | 9.9 | $3.6 \times 10^{-4}$ | $\pm 0.5 \times 10^{-4}$ |
| | | 3 | 97.9 | 2.1 | $4.4 \times 10^{-4}$ | $\pm 0.4 \times 10^{-4}$ |
| MLS 2 | quantity | replicate | % positive | % negative | mean | 95% CI |
| | change in mean of $\Phi$ | 1 | 47.6 | 52.4 | $0.9 \times 10^{-5}$ | $\pm 2.0 \times 10^{-5}$ |
| | | 2 | 47.7 | 52.3 | $0.4 \times 10^{-5}$ | $\pm 1.9 \times 10^{-5}$ |
| | | 3 | 45.4 | 54.6 | $-0.1 \times 10^{-5}$ | $\pm 1.8 \times 10^{-5}$ |
| | colony-level transmission | 1 | 2.5 | 97.5 | $-4.6 \times 10^{-4}$ | $\pm 0.5 \times 10^{-4}$ |
| | | 2 | 4.2 | 95.8 | $-4.8 \times 10^{-4}$ | $\pm 0.3 \times 10^{-4}$ |
| | | 3 | 3.4 | 96.6 | $-4.9 \times 10^{-4}$ | $\pm 0.4 \times 10^{-4}$ |
| | colony-level selection | 1 | 82.6 | 1.6 | $4.7 \times 10^{-4}$ | $\pm 0.4 \times 10^{-4}$ |
| | | 2 | 86.3 | 2.3 | $4.9 \times 10^{-4}$ | $\pm 0.4 \times 10^{-4}$ |
| | | 3 | 82.8 | 3.6 | $4.8 \times 10^{-4}$ | $\pm 0.5 \times 10^{-4}$ |

## Supplementary Movie Captions

**Movie S1**. Movie depicting the dynamics of the simulation described in Fig. 2, which is also shown in Fig. S1 as Replicate 1. Default parameters were used (Table 1). The video plots the positions of all individuals, and the level of altruism of each individual is indicated with the same color scale as in Fig. 2 and 3a. The ticks on the left-hand vertical axis show the scale of altruism, the ticks on the right-hand vertical axis the scale of competition. A high-quality version of this movie is shared here: https://doi.org/10.5281/zenodo.5727313.

**Movie S2**. Movie depicting the dynamics of the simulation Replicate 2 described in Fig. S1. Default parameters were used (Table 1). In the video, the left-hand panel shows the positions of all individuals, as in 5, using the same color scale as in Fig. 2 and 3a. The right-hand panel plots for each position in the habitat the value *A*, which can be interpreted as the amount of public good at that position, as provided by the altruists in the local environment. The ticks on the left-hand vertical axis show the scale of altruism, the ticks on the right-hand vertical axis the scale of competition. A high-quality version of this movie is shared here: https://doi.org/10.5281/zenodo.5727313.

**Movie S3**. Movie depicting the dynamics of the simulation Replicate 3 described in Fig. S1. Default parameters were used (Table 1). In the video, the left-hand panel shows the positions of all individuals, as in 5, using the same color scale as in Fig. 2 and 3a. The right-hand panel plots for each position in the habitat the value *A*, which can be interpreted as the amount of public good at that position, as provided by the altruists in the local environment. The ticks on the left-hand vertical axis show the scale of altruism, the ticks on the right-hand vertical axis the scale of competition. A high-quality version of this movie is shared here: https://doi.org/10.5281/zenodo.5727313.

## References

1. Frank, S. A. Foundations of Social Evolution (Princeton University Press, Princeton, 1998).

2. Marshall, J. A. R. Social Evolution and Inclusive Fitness Theory : An Introduction (Princeton University Press, Princeton, 2015).

3. Nowak, M. A. Five Rules for the Evolution of Cooperation. Science 314, 1560–1563 (Dec. 8, 2006).

4. Cremer, J. et al. Cooperation in Microbial Populations: Theory and Experimental Model Systems. Journal of Molecular Biology. Underlying Mechanisms of Bacterial Phenotypic Heterogeneity and Sociobiology 431, 4599–4644 (Nov. 22, 2019).

5. Kerr, B., Godfrey-Smith, P. & Feldman, M. W. What Is Altruism? Trends in Ecology & Evolution 19, 135–140 (Mar. 1, 2004).

6. West, S. A., Griffin, A. S. & Gardner, A. Social Semantics: Altruism, Cooperation, Mutualism, Strong Reciprocity and Group Selection. Journal of Evolutionary Biology 20, 415–432 (Mar. 2007).

7. Hamilton, W. D. The Evolution of Altruistic Behavior. The American Naturalist 97, 354–356 (Sept. 1, 1963).

8. Sober, P. E., Wilson, P. D. S. & Wilson, D. S. Unto Others: The Evolution and Psychology of Unselfish Behavior New edition. 416 pp. (Harvard University Press, Cambridge, Mass., Oct. 1, 1999).

9. Fletcher, J. A. & Doebeli, M. A Simple and General Explanation for the Evolution of Altruism. Proceedings of the Royal Society of London B: Biological Sciences 276, 13–19 (Jan. 7, 2009).

10. Levin, B. R. & Kilmer, W. L. Interdemic Selection and the Evolution of Altruism: A Computer Simulation Study. Evolution 28, 527–545 (1974).

11. Wilson, D. S. A Theory of Group Selection. Proceedings of the national academy of sciences 72, 143–146 (1975).

12. Wilson, D. S. The Group Selection Controversy: History and Current Status. Annu. Rev. Ecol. Syst. 14, 159–187 (Nov. 1983).

13. Okasha, S. Evolution and the Levels of Selection (Oxford University Press, 2006).

14. Traulsen, A. & Nowak, M. A. Evolution of Cooperation by Multilevel Selection. PNAS 103, 10952–10955 (July 18, 2006).

15. Hogeweg, P. & Takeuchi, N. Multilevel Selection in Models of Prebiotic Evolution: Compartments and Spatial Self-Organization. Orig Life Evol Biosph 33, 375–403 (Oct. 2003).

16. Lehmann, L., Perrin, N. & Rousset, F. Population Demography and the Evolution of Helping Behaviors. Evolution 60, 1137–1151 (2006).

17. Buss, L. W. The Evolution of Individuality 201 pp. (Princeton University Press, 1987).

18. Hauert, C. & Doebeli, M. Spatial Structure Often Inhibits the Evolution of Cooperation in the Snowdrift Game. Nature 428, 643–646 (6983 Apr. 2004).

19. Lion, S. & van Baalen, M. Self-Structuring in Spatial Evolutionary Ecology. Ecology Letters 11, 277–295 (2008).

20. Colizzi, E. S. & Hogeweg, P. High Cost Enhances Cooperation through the Interplay between Evolution and Self-Organisation. BMC Evolutionary Biology 16, 31 (Feb. 1, 2016).

21. Lion, S. & Gandon, S. Habitat Saturation and the Spatial Evolutionary Ecology of Altruism. Journal of Evolutionary Biology 22, 1487–1502 (2009).

22. Wakano, J. Y., Nowak, M. A. & Hauert, C. Spatial Dynamics of Ecological Public Goods. PNAS 106, 7910–7914 (May 12, 2009).

23. Hamilton, W. D. The Genetical Evolution of Social Behaviour. I. Journal of theoretical biology 7, 1–16 (1964).

24. Grafen, A. A Geometric View of Relatedness. Oxford surveys in evolutionary biology 2, 28–89 (1985).

25. Nowak, M. A. & May, R. M. Evolutionary Games and Spatial Chaos. Nature 359, 826–829 (1992).

26. Taylor P. D. Altruism in Viscous Populations — an Inclusive Fitness Model. Evol Ecol 6, 352–356 (July 1, 1992).

27. Wilson, D. S., Pollock, G. B. & Dugatkin, L. A. Can Altruism Evolve in Purely Viscous Populations? Evol Ecol 6, 331–341 (July 1, 1992).

28. Queller, D. C. Does Population Viscosity Promote Kin Selection? Trends in Ecology & Evolution 7, 322–324 (Oct. 1, 1992).

29. West, S. A. Cooperation and Competition Between Relatives. Science 296, 72–75 (Apr. 5, 2002).

30. Wallace, B. Hard and Soft Selection Revisited. Evolution 29, 465–473 (1975).

31. Goodnight, C. J., Schwartz, J. M. & Stevens, L. Contextual Analysis of Models of Group Selection, Soft Selection, Hard Selection, and the Evolution of Altruism. The American Naturalist 140, 743–761 (1992).

32. Mitteldorf, J. & Wilson, D. S. Population Viscosity and the Evolution of Altruism. Journal of Theoretical Biology 204, 481–496 (June 2000).

33. Queller, D. C. Genetic Relatedness in Viscous Populations. Evolutionary Ecology 8, 70–73 (Jan. 1994).

34. Wilson, R. L., Urbanowski, M. L. & Stauffer, G. V. DNA Binding Sites of the LysR-type Regulator GcvA in the Gcv and gcvA Control Regions of Escherichia Coli. Journal of bacteriology 177, 4940–4946 (1995).

35. Chuang, J. S., Rivoire, O. & Leibler, S. Simpson’s Paradox in a Synthetic Microbial System. Science 323, 272–275 (Jan. 9, 2009).

36. Cremer, J., Melbinger, A. & Frey, E. Growth Dynamics and the Evolution of Cooperation in Microbial Populations. Scientific Reports 2 (Feb. 21, 2012).

37. Slatkin, M. Gene Flow and Genetic Drift in a Species Subject to Frequent Local Extinctions. Theoretical Population Biology 12, 253–262 (Dec. 1, 1977).

38. Hardin, G. The Tragedy of the Commons. Science 162, 1243–1248 (Dec. 13, 1968).

39. Turing, A. M. The Chemical Basis of Morphogenesis. Philosophical Transactions of the Royal Society of London. Series B, Biological Sciences 237, 37–72 (1952).

40. Price, G. R. Extension of Covariance Selection Mathematics. Ann. Hum. Genet. 35, 485–490 (Apr. 1972).

41. Damuth, J. & Heisler, I. L. Alternative Formulations of Multilevel Selection. Biol Philos 3, 407–430 (Oct. 1, 1988).

42. Price, G. R. Selection and Covariance. Nature 227, 520–521 (Aug. 1, 1970).

43. Frank, S. A. Natural Selection. IV. The Price Equation. J Evol Biol 25, 1002–1019 (June 2012).

44. Maynard Smith, J. in The Latest and the Best: Essays on Evolution, Ed Dupré J. 119–131 (1987).

45. Gardner, A. & Grafen, A. Capturing the Superorganism: A Formal Theory of Group Adaptation. Journal of Evolutionary Biology 22, 659–671 (2009).

46. Black, A. J., Bourrat, P. & Rainey, P. B. Ecological Scaffolding and the Evolution of Individuality. Nat Ecol Evol 4, 426–436 (3 Mar. 2020).

47. Wilson, D. S. Structured Demes and Trait-Group Variation. The American Naturalist 113, 606–610 (Apr. 1979).

48. Powers, S. T., Penn, A. S. & Watson, R. A. The Concurrent Evolution of Cooperation and the Population Structures That Support It. Evolution 65, 1527–1543 (June 2011).

49. Boerlijst, M. C. & Hogeweg, P. Spiral Wave Structure in Pre-Biotic Evolution: Hypercycles Stable against Parasites. Physica D: Nonlinear Phenomena 48, 17–28 (Feb. 1, 1991).

50. De Jager, M., Weissing, F. J. & van de Koppel, J. Why Mussels Stick Together: Spatial Self-Organization Affects the Evolution of Cooperation. Evol Ecol 31, 547–558 (Aug. 2017).

51. Van Ballegooijen, W. M. & Boerlijst, M. C. Emergent Trade-Offs and Selection for Outbreak Frequency in Spatial Epidemics. Proc Natl Acad Sci U S A 101, 18246–18250 (Dec. 28, 2004).

52. Takeuchi, N. & Hogeweg, P. Multilevel Selection in Models of Prebiotic Evolution II: A Direct Comparison of Compartmentalization and Spatial Self-Organization. PLoS Comput Biol 5, e1000542 (Oct. 2009).

53. Rietkerk, M. & van de Koppel, J. Regular Pattern Formation in Real Ecosystems. Trends in Ecology & Evolution 23, 169–175 (Mar. 1, 2008).

54. Lewontin, R. C. The Units of Selection. Annual Review of Ecology and Systematics, 1–18 (1970).

55. Okasha, S. Maynard Smith on the Levels of Selection Question. Biol Philos 20, 989–1010 (Nov. 1, 2005).

56. Goodnight, K. F. The Effect of Stochastic Variation on Kin Selection in a Budding-Viscous Population. The American Naturalist 140, 1028–1040 (Dec. 1992).

57. Gardner, A. & West, S. A. Demography, Altruism, and the Benefits of Budding. Journal of Evolutionary Biology 19, 1707–1716 (2006).

58. Henriques, G. J. B., van Vliet, S. & Doebeli, M. Multilevel Selection Favors Fragmentation Modes That Maintain Cooperative Interactions in Multispecies Communities. PLoS Comput Biol 17 (ed Traulsen, A.) e1008896 (Sept. 13, 2021).

59. Nowak, M. A., Tarnita, C. E. & Wilson, E. O. The Evolution of Eusociality. Nature 466, 1057–1062 (Aug. 26, 2010).

60. Abbot, P. et al. Inclusive Fitness Theory and Eusociality. Nature 471, E1–E4 (7339 Mar. 2011).

61. Strassmann, J. E., Page, R. E., Robinson, G. E. & Seeley, T. D. Kin Selection and Eusociality. Nature 471, E5–E6 (7339 Mar. 2011).

62. Queller, D. C. Kin Selection and Its Discontents. Philosophy of Science 83, 861–872 (Dec. 2016).

63. Levin, S. R. & Grafen, A. Inclusive Fitness Is an Indispensable Approximation for Understanding Organismal Design. Evolution 73, 1066–1076 (2019).

64. Goodnight, C. J. Contextual Analysis and Group Selection. Behavioral and Brain Sciences 17, 622 (1994).

65. Doekes, H. M. & Hermsen, R. Multiscale Selection in Spatially Structured PopulationsDec. 21, 2021.

66. Rice, S. H. Evolutionary Theory: Mathematical and Conceptual Foundations 370 pp. (Sinauer Associates, 2004).

67. Waters, C. K. Okasha’s Unintended Argument for Toolbox Theorizing. Philosophy and Phenomenological Research 82, 232–240 (2011).

68. Queller, D. C. A General Model for Kin Selection. Evolution 46, 376–380 (1992).

69. Flyvbjerg, H. & Petersen, H. G. Error Estimates on Averages of Correlated Data. The Journal of Chemical Physics 91, 461–466 (July 1989).

70. Brandon, R. N. Adaptation and Evolutionary Theory. Studies in History and Philosophy of Science Part A 9, 181–206 (Sept. 1, 1978).

71. Mills, S. K. & Beatty, J. H. The Propensity Interpretation of Fitness. Philosophy of Science 46, 263–286 (June 1979).

